# Cryo-OrbiSIMS for 3D molecular imaging of a bacterial biofilm in its native state

**DOI:** 10.1101/859967

**Authors:** Junting Zhang, James Brown, David Scurr, Anwen Bullen, Kirsty MacLellan-Gibson, Paul Williams, Morgan R. Alexander, Kim R. Hardie, Ian S. Gilmore, Paulina D. Rakowska

## Abstract

We describe a method for label-free molecular imaging of biological materials, preserved in a native state, by using an OrbiSIMS instrument equipped with cryogenic sample handling and developing a high-pressure freezing protocol compatible with mass spectrometry. We studied the 3D distribution of quorum sensing signalling molecules, nucleobases and bacterial membrane molecules, in a mature *Pseudomonas aeruginosa* biofilm, with high spatial-resolution and high mass-resolution.

## Main

Recent progress in mass spectrometry imaging methods, including matrix assisted laser desorption/ionisation mass spectrometry (MALDI MS) and secondary ion mass spectrometry (SIMS), enables label-free molecular imaging with sub-cellular resolution.^1, 2, 3, 4^ We previously introduced the 3D OrbiSIMS,^3^ a hybrid instrument with a time-of-flight (ToF) mass spectrometer (MS) for high-speed 3D imaging and a high-field Orbitrap MS (mass resolving power of 240,000 at *m*/*z* 200 and a mass accuracy of < 2 p.p.m) for 2D imaging and molecular identification. An argon gas cluster ion beam enables imaging of biomolecules with a spatial resolution of < 2 μm with low fragmentation.^3^

Unfortunately, the SIMS high-vacuum operating conditions (< 10^-8^ mbar) necessitate special biological sample preparation. A popular protocol is fixation (chemical or cryogenic) followed by drying.^5^ Although morphology is preserved in some cases, the biggest risk of these protocols is chemical redistribution artefacts, such as migration of cholesterol in brain tissue to the surface^6^ or redistribution of lipids in adipose tissue during OsO_4_ staining.^7^ Alternatively, a cryogenic sample holder allows analysis in the frozen-hydrated state after cryo-fixation followed either by fracturing in vacuum or by direct transfer to the instrument. The method was shown to prevent the chemical redistribution.^8,9,10^

Cryogenic sample preparation techniques need to avoid water crystal formation in the sample, which could cause the rupture of cellular membranes and the movement of molecules. Plunge-freezing is the most popular method used for SIMS studies of cells. Whilst cell morphology can be preserved, the freezing rates are only sufficiently fast to achieve vitreous ice in thin samples (< 10-20 μm). Plunge freezing is not suitable for thicker samples such as tissues due to formation of crystalline ice.^11^ More sophisticated sample preparation protocols were developed by Nygren et al.^12^ and Magnusson et al.^13^ over a decade ago, using high-pressure freezing followed by freeze-fracture and then freeze drying for ToF-SIMS imaging at room temperature. They demonstrated that muscle tissue structure was well preserved and analytes, such as Na^+^ and K^+^, were retained at their original locations.

Here, we report the development of cryo-OrbiSIMS for analysis of biological samples preserved in their native state. The instrument is equipped with a cryo-stage and a docking station, compatible with a cryogenic sample transfer shuttle used extensively in electron microscopy. This provides opportunities for future correlative imaging experiments. The cryo-OrbiSIMS workflow is presented in **Fig. 1**. The sample is prepared on a suitable substrate for high-pressure freezing (HPF) (**Fig. 1a**) and then assembled into a sandwich with specimen carriers for HPF (**Fig. 1b**). A cryo-protectant or filler is used (**Fig. 1b**). HPF simultaneously applies a pressure of 210 kPa and cooling to a temperature of −196 °C (**Fig. 1c**). Samples are mounted on to a specimen holder under liquid nitrogen in a loading station and transferred under the vapour phase of liquid nitrogen into the pre-cooled cryogenic sample transfer shuttle (**Fig. 1d**). The sample temperature is kept at below −150 °C. Finally, the shuttle is docked with the OrbiSIMS (**Fig 1e**) where the sample is transferred under vacuum into a pre-cooled parking position in the load lock. This approach overcomes the problem of water condensation on the sample surface reported in previous cryo-SIMS studies.^14^ A SIMS depth profile through a frozen-hydrated bacterial biofilm using a 30 keV Bi_3_^+^ beam and TOF-MS for analysis with Ar_3000_^+^ for sputtering (**Supplementary Fig. 1**) shows no initial high water signal (H_3_O^+^) from an adventitious ice layer is evident.

**Figure 1.**
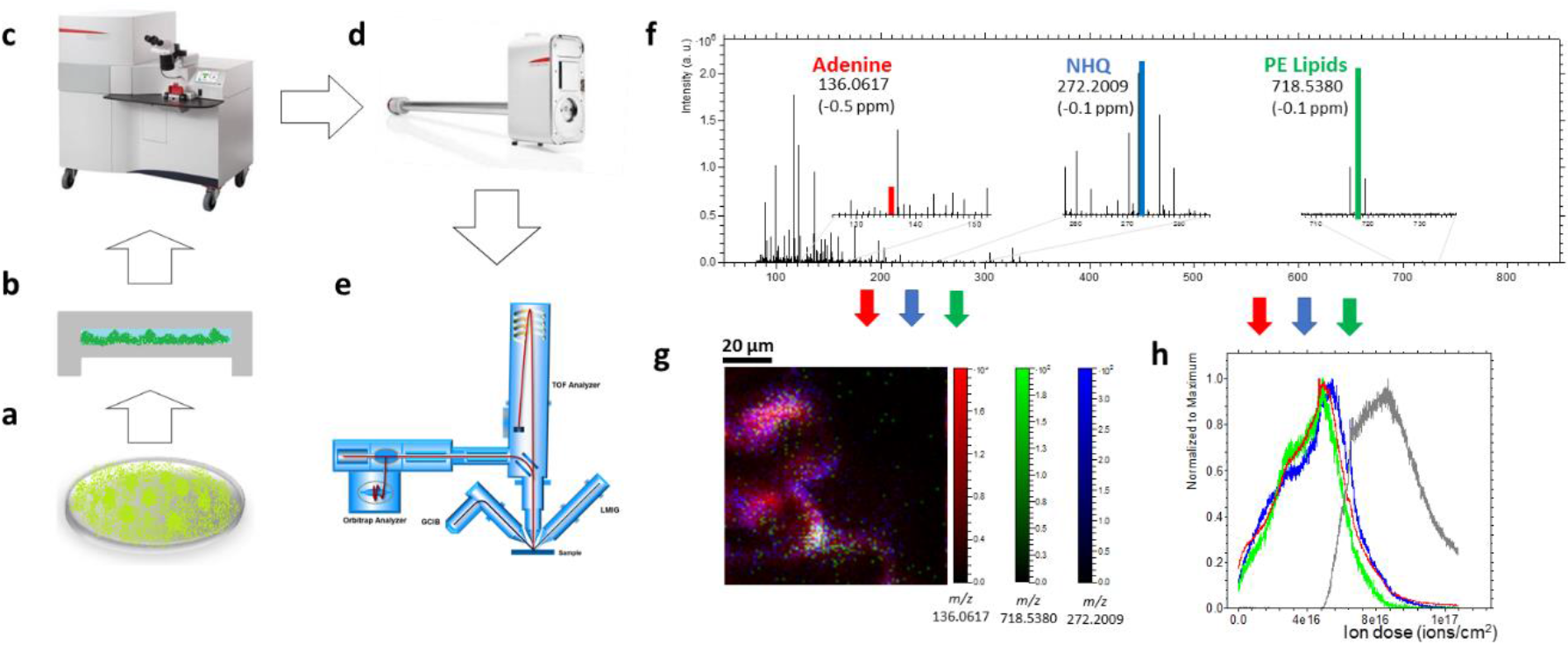
The workflow of sample preparation and cryo-OrbiSIMS analysis of biofilms. (a-e) Schematic drawings showing the experimental procedures: (a) Biofilm growing on the Al sample carrier, (b) specimen carrier assembly with ammonium formate added as a cryo-protectant, (c) high pressure freezing, (d) sample transfer to the 3D OrbiSIMS instrument using the vacuum cryo transfer, (e) analysis of samples by cryo-OrbiSIMS, (f) 20 keV Ar_3000_^+^ Orbitrap MS spectrum of a frozen-hydrated *P. aeruginosa* biofilm. The spectrum is the sum of 150 scans with a total ion dose 3.4×10^16^ ions/cm^2^. The *m/z* values used for RGB MS image (g) and depth profile (h) are color-coded: (g) MS image of adenine at *m/z* 136.0617 (red), NHQ at *m/z* 272.2009 (blue), PE lipids at *m/z* 718.5380 (green as an RGB overlay), (h) intensity depth profile normalised to the maximum intensity of adenine at *m/z* 136.0618 (red line), NHQ at *m/z* 272.2007 (blue line), PE lipids at *m/z* 718.5380 (green line) and Al_2_O_3_(H_2_O)_3_H^+^ at m/z 156.9866 (grey line).

We demonstrate the capability of the cryo-OrbiSIMS by studying a frozen-hydrated mature biofilm of *P. aeruginosa* (strain PAO1), which is an important, well-studied human pathogen.^15^ Biofilms are structured communities of bacteria in a self-produced extracellular matrix^16^ consisting mostly of water (90%)^17^ and containing extracellular DNA, exopolysaccharides, lipids and proteins. Biofilms contain also quorum sensing (QS) signalling molecules used for bacterial cell-cell communication and involved in biofilm maturation.^18^

The heterogenous distribution of microbial cells and their metabolites within a biofilm requires analytical techniques capable of spatial mapping of biochemical molecules. Confocal laser scanning microscopy (CLSM), in combination with fluorescent labels, is a popular technique to study biofilms^19^. However, it requires labelling and provides limited information on QS signalling molecules. SIMS is a complementary technique, previously used to image quinolones, surfactants and antibiotics in biofilms^20,21^ and microbial colonies.^22^ However, in these studies the biofilms were dried before analysis. The only report for SIMS analysis of a hydrated biofilm is at the unstable liquid-vacuum interface of an *in vacuo* exposed biofilm, described by Ding et al.^23^ Their methodology gives a small field of view (2 μm diameter) and limited molecular information.^23^

Sample preparation for bacterial biofilm by high-pressure freezing is key to retaining the hydrated structure of a biofilm as it consists of over 90% water and the average thickness of mature biofilm always more than 20 μm.^24^ Bacterial biofilm is directly grown on the aluminium (Al) specimen carriers, which are normally used for HPF. Biofilm could grow well on this substrate which is shown in CLSM (**Supplementary Fig. 6**), there is also added advantage of the Al substrate conductivity, which helps reducing the charging issue in SIMS. A cryo-protectant is often employed in high-pressure freezing to fill the residual gap between the sample and the specimen carriers, otherwise freezing damage can occur. A wide range of cryo-protectants are used in cryo-EM.^25^ We evaluated the compatibility of common EM cryoprotectants, including dextran, 1-hexadecane, bovine serum albumin (BSA), and methanol for suitability with SIMS (see on-line methods) by using pellets of *Escherichia coli*. All of these cryoprotectants suppressed the signal intensity of biomolecular ions (**Supplementary Fig. 2**). We found that ammonium formate, often used to remove salts prior to SIMS analysis,^5^ could also be used as a cryo-protectant. Analysis by cryo-EM showed bacterial cells are preserved very well and that the material is vitreous as determined by electron diffraction (**Supplementary Fig. 3**). This method did not produce any interfering signals in the Orbitrap mass spectrum and signals from lipids and various metabolites could be detected (**Supplementary Fig. 2**).

A cryo-OrbiSIMS image (**Supplementary protocol**) with Orbitrap MS image of a frozen-hydrated *P. aeruginosa* biofilm (**Fig. 1f**) shows the spatial distribution of adenine (C_5_H_6_N_5_^+^, [M+H]^+^, *m*/*z* 136.0617, red) (**Supplementary Fig. 5**), 2-nonyl-4-hydroxyquinoline (NHQ; C_18_H_26_NO^+^, [M+H]^+^, *m*/*z* 272.2007, blue) and a phosphatidylethanolamine (PE) lipid (C_39_H_77_NO_8_P^+^, [M+H]^+^, *m*/*z* 718.5381, green). From the average mass spectrum of depth profile (**Fig. 1f**) we annotate 87 different compounds including 5 nucleobases, 20 amino acids, 2 PE lipids, 33 alkyl quinolones and 6 *N*-acyl homoserine lactones (QS signalling molecules) with a mass error < 2 ppm. (**Supplementary Table 1**). MS/MS analysis of six alkyl quinolones *m*/*z* 244.1694, *m*/*z* 260.1643, *m*/*z* 270.1854, *m*/*z* 272.2010, *m*/*z* 286.1799, *m*/*z* 288.1959 support their annotation as HHQ [M+H]^+^, HQNO/C7-PQS [M+H]^+^, C9:1-AQ [M+H]^+^, NHQ [M+H]^+^, C9:1-NO [M+H]^+^ and C9-PQS/NQNO [M+H]^+^ (**Supplementary Fig. 4**). A single beam 20 keV Ar_3000_^+^ Orbitrap MS depth profile (**Fig. 1h**) for the same ions in the image (**Fig. 1f**) shows their variation in concentration throughout the depth of the entire biofilm to the aluminium substrate (Al_2_O_3_H_7_O_3_^+^, *m*/*z* 156.9866, grey line). Interestingly, different alkyl quinolone signals show slightly different trends in the depth profile (**Supplementary Fig. 6**), suggesting different vertical distributions.

Since biofilms have complex architectures, it is important to study them in 3D. However, Orbitrap MS imaging with high mass resolution is slow, and therefore impractical to render 3D images with high definition in the vertical direction. Here, we only showed eight subsequent Orbitrap MS images (summed in pairs) with 100 × 100 μm FoV (field of view) in the area close to the bottom, which was randomly selected as an example. Adenine (C_5_H_6_N_5_^+^, [M+H]^+^, *m*/*z* 136.0617) is a nucleic acid marker and can originate from both the cytoplasm of the bacteria and the extracellular DNA present in the extracellular matrix, whilst the PE lipid headgroup (C_2_H_9_PNO_4_^+^, [M+H]^+^, *m/z* 142.0262) is a marker for the bacterial membrane and is only associated with bacterial cells and microvesicles. It is notable that one PE lipid molecular ion ((C_39_H_77_NO_8_P^+^, [M+H]^+^, *m*/*z* 718.5381) shows a different distribution to the PE lipid headgroup ion (C_2_H_9_PNO_4_^+^, [M+H]^+^, *m*/*z* 142.0262) (**Supplementary Fig. 9** and **Fig. 10a**) (Pearson correlation coefficient (PCC) = 0.53), suggesting there might be low signal intensity of these PE lipids and that other PE lipids were not detected as intact molecular ions.^26^ Therefore, we use the PE lipid headgroup as a marker for bacterial membranes. The adenine signal correlates highly to the signal of PE lipid headgroup ion (**Fig. 2a-b** and **Supplementary Fig. 10c**) (PCC = 0.95), suggesting that the detected adenine comes mostly from bacterial cells. It might be that the concentration of extracellular DNA is too low in the biofilms for the adenine to be detected by SIMS. The distribution of NHQ is similar to that of adenine (PCC = 0.80) and the PE lipid headgroup (PCC=0.75) (**Fig. 2a-c** and **Supplementary Fig. 10b, d**), which suggests that NHQ is co-localized with bacterial cells within the biofilm. NHQ is an extracellular signalling molecule but, because of its physical properties, its high proportion is associated with the cell envelope and any microvesicles that had been shed into the biofilm matrix.^27, 28^ Interestingly, all the signals have a different distribution scan by scan, which might point to the heterogeneity of the biofilm. The distribution of 47 other small molecules in the biofilm was also mapped (**Supplementary Fig. 11**).

**Figure 2.**
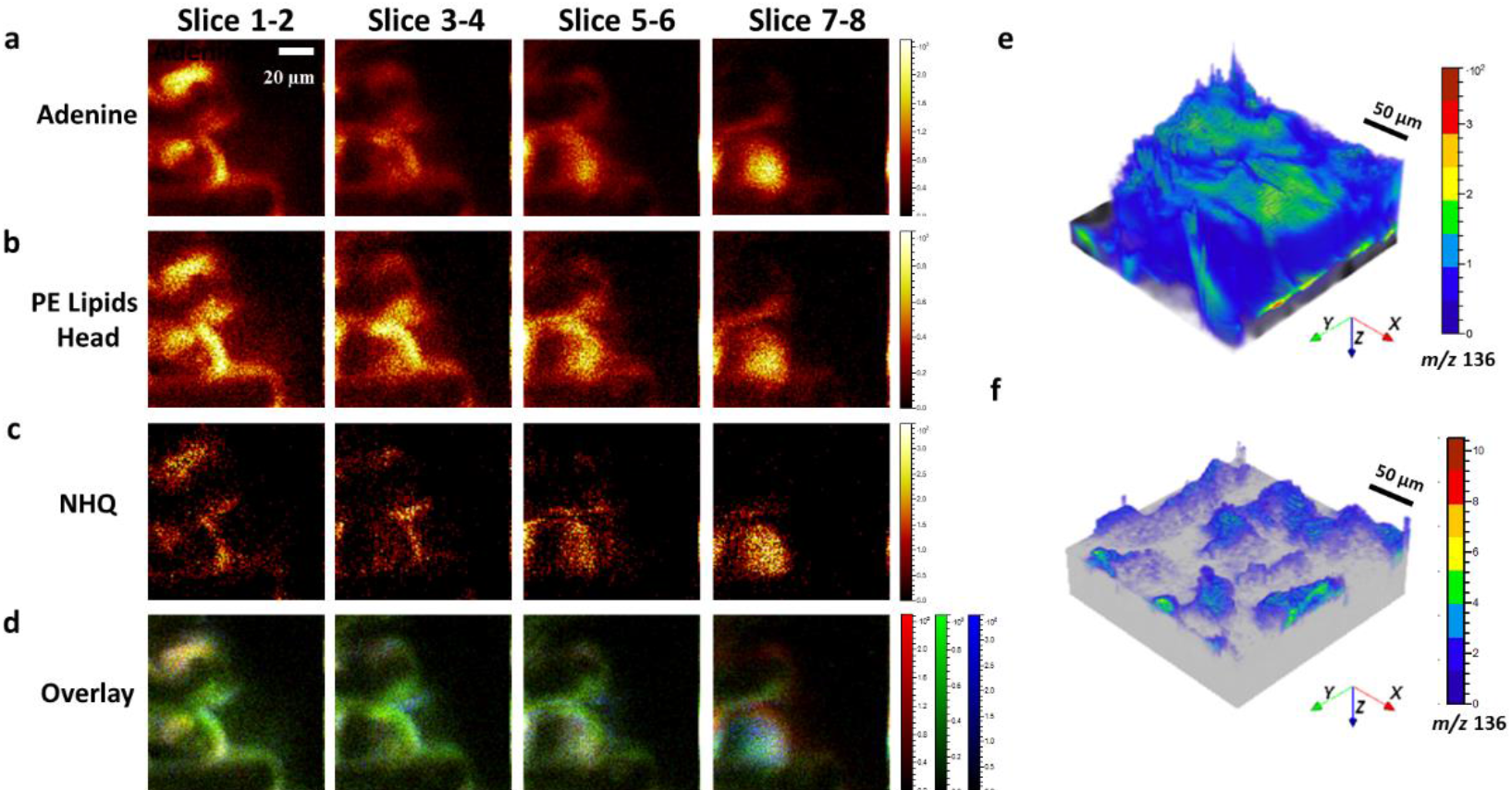
OrbiSIMS images of a frozen-hydrated *P. aeruginosa* biofilm. (a-d) Sequence of 20 keV Ar_3000_^+^ (pixel size 1 μm) Orbitrap MS images of (a) adenine at *m/z* 136.0617, (b) PE lipid headgroups at *m/z* 142.0262, (c) NHQ quinolone at *m/z* 272.2007, (d) RGB overlay of adenine at m/z 136.0617 (red), NHQ at *m/z* 272.2007 (blue), PE lipids head at m/z 142.0262 (green) of frozen-hydrated biofilm at various sputter depths. (e) 3D rendered ToF MS images of the frozen hydrated biofilm with adenine/2-Aminoacetophenone (*m/z* 136) on the substrate (Al, *m/z* 27). (f) 3D rendered ToF MS images of the freeze-dried biofilm with adenine/2-Aminoacetophenone (*m/z* 136) on the substrate (Al, *m/z* 27). The images in (e) and (f) were corrected assuming a flat substrate.

The OrbiSIMS has a unique ability to operate in a dual analyser-dual beam mode, where 3D images are acquired with the high resolution 30 keV Bi_3_^+^ primary ion beam and ToF MS while simultaneously an Orbitrap MS depth profile is acquired during the 20 keV Ar_3000_^+^ sputtering cycles.^3^ As ToF MS imaging is fast, it is possible to generate 3D imaging with a surface correction.^29^ Alkyl quinolone and lipid molecules are not detected in the ToF MS spectra **(Supplementary Fig. 8)**, likely due to much higher fragmentation. However, K^+^ and adenine (C_5_H_6_N_5_^+^, [M+H]^+^, *m*/*z* 136) were detected as markers for bacterial cells, enabling 3D imaging of the bacterial distribution within a biofilm. The Orbitrap data shows that the broad ToF MS peak at *m*/*z* 136 is mainly composed of two peaks, adenine (m/z 136.0617) and 2-aminoacetophenone (*m*/*z* 136.0756), but with the former being the most dominant (> 80%) (**Supplementary Fig. 12**). The latter is a volatile signal molecule derived via a side reaction of the alkylquinolone biosynthetic pathway.^30^ The reconstructed 3D ToF MS image of adenine (C_5_H_6_N_5_^+^, [M+H]^+^, *m*/*z* 136) and K^+^, with a surface correction algorithm defining the aluminium substrate as flat, show the outline of two microcolonies inside the biofilm (**Fig. 2e** and **Supplementary Fig. 13 and Video 1-2**).

Almost all previously published SIMS studies of biofilms concentrated in the analysis of dried samples. Therefore, we made a comparison between data acquired from frozen-hydrated samples to measurements collected from freeze-dried biofilms. For comparison, the sample was re-imaged in a different area following *in situ* freeze drying (**Supplementary Fig. 14** and **Fig. 2f**) and with a corresponding depth profile (**Supplementary Fig. 15**), which shows the biofilm has collapsed after freeze-drying. Interestingly, the quinolone signal for the NHQ (C_18_H_26_NO^+^, *m*/*z* 272) ion becomes detectable in the 30 keV Bi_3_^+^ ToF MS (**Supplementary Fig. 16**), presumably due to the higher concentration of molecules within the analysed volume. From the 20 keV Ar_3000_^+^ Orbitrap data we measured the total signal intensity (methods) for 87 nucleobases, amino acids, alkyl quinolones, lipids and other molecules and the intensity ratio for frozen-hydrated to freeze-dried (**Supplementary Fig. 17**) sample preparation (**Supplementary Table 2**). All intensities are enhanced in the frozen-hydrated biofilm probably because the low gas phase basicity (665 kJ/mol) of the water matrix facilitates proton transfer for ionisation.^31,32^ Water can also stabilize and trap both the cation and anion to reduce the ion suppression by salts.^33^ We generally found that polar molecules (low Log *P*) are more strongly enhanced with amino acid ions typically 10,000 fold higher (**Supplementary Fig. 18**).

In conclusion, we report a method for chemical imaging of biological samples in their native state using a cryo-OrbiSIMS with a protocol for sample preparation and handling. We show chemical mapping of a frozen-hydrated biofilm in 2D and 3D. Analysis with the 20 keV Ar_3000_^+^ and Orbitrap MS gives significantly more biological information compared with 30 keV Bi_3_^+^ and ToF MS. High-pressure freezing is critical to create vitreous ice and preserve the biofilm native architecture. We also found that the SIMS sensitivity of polar molecules is increased by more than 10,000-fold in the hydrated state. Our method can easily be adapted to various other biological systems such as tissues and cells and is compatible with cryo-EM for correlative imaging.

## Acknowledgements

The authors thank Julie Watts, Nigel Halliday, Mike Shaw, Helen Lewis, Jean-Luc Vorng and Robert Francis for their technical support during the project. The authors thank M. Tiddia for laser microscopy measurement of the HPF specimen carriers. This work forms part of the 3D OrbiSIMS project in the Life-science and Health programme of the National Measurement System of the UK Department of Business, Energy and Industrial strategy. This work has received funding from the MetVBadBugs project of the EMPIR programme co-financed by the Participating States and from the European Union’s Horizon 2020 research and innovation programme. The EPSRC are acknowledged for the award of a Strategic Equipment grant to the University of Nottingham for the 3DOrbisSIMS facility (EP/P029868/1).

## Author contributions

All authors contributed to and approved the manuscript. JZ, DS and PDR acquired OrbiSIMS and ToF-SIMS data. JB developed method and grew mature biofilms and performed confocal microscopy. KH, PW and JB designed biofilm experiments. JZ, PDR, KMG, AW, ISG developed HPF and cryo-analysis protocol. ISG created OrbiSIMS concept and cryo sample handling design concept with PDR. KMG and AB performed cryo-EM analysis. JZ, PDR and ISG designed OrbiSIMS experiments. JZ analysed and interpreted SIMS data with additional interpretation from PDR, ISG, MRA and DS. MRA oversaw OrbiSIMS experiments at Nottingham. JZ, ISG, PDR and MRA wrote the paper.

## Methods

### Sample preparations

*Pseudomonas aeruginosa* strains used in this experiment were PAO1, (Holloway, B. W. 1955. Genetic recombination in *Pseudomonas aeruginosa*. J. Gen. Microbiol. 13:572-581) and PAO1 transformed with the red fluorescent protein mCherry regulated by a constitutive promoter (pME6032-ptac::mCherry). Strains were maintained on Lysogeny agar and grown overnight in Lysogeny Broth (LB) at 37°C with constant shaking. For growth of biofilms *P. aeruginosa* grown overnight were diluted to an OD_600_ =0.05 in FAB medium [2 g (NH_4_)2SO_4_, 6 g Na_2_HPO_4_ · 2H_2_O, 3 g KH_2_PO_4_, 3 g NaCl per litre] with 0.1 mM CaCl2, 1 mM MgCl_2_, 1 ml.L^-1^ trace metals mix (200 mg.L^-1^ CaSO_4_ · 2H_2_O, 200 mg.L^-1^ FeSO_4_ · 7H_2_O, 20.L^-1^ MnSO_4_ · H_2_O, 20 mg.L^-1^ CuSO_4_ · 5H_2_O, 20 mg.L^-1^ ZnSO_4_ · 7H_2_O, 10 mg.L^-1^ CoSO_4_ · 7H_2_O, 12 mg.L^-1^ NaMoO_4_ · H_2_O, and 5 mg.L^-1^ H_3_BO_3_) and 30 mM glucose. Biofilms were directly grown on 3 mm aluminium sample carriers flat face over 48 h using a rotary flow system. Growth medium was replaced after 24 h. The fresh biofilms were washed 2-3 times with 150 mM ammonium formate solution and assembled in a sample carrier system for high pressure freezing (detailed protocols are shown in the **Supplementary Protocols**). After high pressure freezing, samples were stored in liquid nitrogen. The *Escherichia coli* (*E. coli*) used in the cryo-protectant selection experiments was maintained on Lysogeny agar and grown overnight in LB (LB) at 37°C with constant shaking. For preparing *E. coli* pellets for HPF, the cells were washed with 150 mM ammonium formate 2-3 times and gently pelleted by centrifugation (3,000 rpm maximum). The cells were resuspended in the cryo-protectant (20% dextran, 1-hexadecane, 5% bovine serum albumin (BSA), 20% methanol and 150 mM ammonium formate). Approximately 1 μl of the cell suspension was pipetted into a well of an Al specimen carrier and assembled as a sandwich for HPF. For the analysis of the freeze-dried biofilm, the sample was freeze-dried in the load lock of OrbiSIMS allowing the sample to slowly warm to room temperature over a period of 12 h.

### Cryo-OrbiSIMS experimental methods

The cryo-OrbiSIMS is equipped with a fully proportional–integral–derivative (PID) temperature controller which controls resistive heating and a direct liquid nitrogen (LN_2_) closed loop circulation cooling stage, allowing sample temperature control within the load lock and main chamber. Being installed with cryogenic storage tanks, LN_2_ was pumped for circulating the cooling media through vacuum feed-throughs to a cooling finger below the sample, allowing fast cooling to −180 °C with a stability of ± 1-2 °C for at least 7 days. This system allows for full sample movement in *x*, *y*, *z*, rotate and tilt directions whilst under cryogenic conditions.

The protocols for transferring samples to Cryo-OrbiSIMS, setting up Cryo-3D OrbiSIMS and loading samples are provided in the **Supplementary Protocols**. The data shown in **Supplementary Fig. 4, 14 and 16** were obtained from the NPL 3D OrbiSIMS (Hybrid SIMS, IONTOF GmbH, Germany) and the remaining data were obtained from the University of Nottingham 3D OrbiSIMS (Hybrid SIMS, IONTOF GmbH, Germany). All cryo-OrbiSIMS analyses (except **Supplementary Fig. 4**) were conducted at −180 °C. Mass calibration of the Q Exactive instrument was performed once a day using silver cluster ions. Electrons with an energy of 21 eV and a current of 10 μA, and argon gas flooding were used for charge compensation. Three modes of 3D OrbiSIMS were mainly used for the work described in this paper (Details on the operation mode are given in ref. 3): mode 4 (single beam, 20 keV Ar_3000_^+^, Orbitrap MS) including **Fig. 1f** and **1h**, **Supplementary Fig. 2, 7, 15** and **17**; mode 10 (dual beam dual analyser depth profile, 30 keV Bi_3_^+^ with ToF MS imaging, 20 keV Ar_3000_^+^ with Orbitrap MS) including **Fig. 2e** and **f**, **Supplementary Fig. 1, 8, 12, 13, 14** and **16**; mode 8 (single beam, 20 keV Ar_3000_^+^, Orbitrap MS imaging) including **Fig. 1g, 2a-d**, **Supplementary Fig. 5, 9, 11**. For all Orbitrap data, mass spectral information was collected from a mass range from 80-1200 Da. The Orbitrap analyser was operated in positive-ion mode at the 240,000 at m/z 200 mass-resolution setting (512 ms transient time). For depth profiling as in **Fig. 1h**, **Supplementary Fig. 7** and **15**, a 300 × 300 μm region of sample was sputtered using defocused 20 keV Ar_3000_^+^ beam for 2.0 s per cycle. The total ion dose was 1.17-1.27×10^17^ ions/cm^2^. For ToF 3D imaging in **Fig. 2e** and **f**, **Supplementary Fig. 13** and **14**, 300 × 300 μm images with 256 × 256 pixels were obtained using a pulsed 30 keV Bi_3_^+^ with a beam current of 0.15 pA with ToF MS analyser. These images are interleaved with Orbitrap MS acquired during 20keV Ar_3000_^+^ GCIB sputtering cycle from same field of view, but with an additional sputter border of 20 μm width to avoid edge effects. In **Fig. 2e** and **Supplementary Fig. 13**, 1760 scans were accumulated in the ToF images, which correspond to a total primary ion dose of 1.60×10^14^ ions/cm^2^. The total GCIB sputtering dose was 4.00 × 10^17^ ions/cm^2^. For **Fig. 2f** and **Supplementary Fig. 14**, 395 scans were accumulated for the ToF images, which correspond to a total primary ion dose of 4.04 × 10^13^ ions/cm^2^. The total GCIB sputtering dose was 8.08 × 10^17^ ions/cm^2^. For Orbitrap 2D and 3D imaging in **Fig. 2a-d** and **Supplementary Fig.5**, **9** and **11**. images containing 100 × 100 pixels were acquired over an area of 100 × 100 μm. Approximately 2500 shots at 200 μs per cycle were accumulated in the C-trap per pixel. The total ion dose was 4.21×10^16^ ions/cm^2^.

### Confocal Microscopy experimental methods

A Zeiss LSM 700 microscopy was used to image the biofilms and Zeiss Zen software was used for image processing. Cells were visualised via expression of the red fluorescent protein mCherry. Extracellular DNA in the biofilm was visualised using the dye YOYO-1 (0.1 μM; ThermoFisher). YOYO-1 was allowed to interact with the biofilm for 15 min prior to microscopy.

### Data Analysis

The 3D OrbiSIMS was controlled by software provided by SurfaceLab Version 7.0 (ION-TOF, Germany), which used the application programming interface (API) provided by Thermo Fisher for both control of the Orbitrap MS portion of the instrument as well as online retrieval of the data. Both ToF-SIMS and Orbitrap MS image analyses were performed using SurfaceLab Version 7.0 (ION-TOF, Germany). In **Fig. 2e, f, Supplementary Fig. 13** and **14**, the ToF-SIMS ion images for *m*/*z* 39, 136 and 272 were corrected with shift correction. 3D renderings were constructed using a so-called volume visualization with the *xy* binning of 16 pixels and *z* binning of 16 scans same binning using SurfaceLab Version 7. The 3D image was created after vertical shift correction (SurfaceLab Version 7), which means *z* position of the voxels was adjusted to take the initial sample topography into account, under the assumption that the aluminium substrate is a uniform flat plane. The image correlation calculations were performed in GraphPad Prism8 using a Pearson’s (two-tailed) correlation. test at the 95% confidence level. The comparison of frozen-hydrated and dehydrated biofilm was from the 20 keV Ar_3000_^+^ Orbitrap MS depth profile data by integrating the mass spectrum from the surface until the aluminium substrate marker ion (Al_2_O_3_(H_2_O)3H^+^ at *m/z* 156.9866 in the frozen sample and Al_7_^+^ at *m/z* 188.8702 in the dehydrated sample) reached 90 % of its maximum intensity.

