## Supplementary Information for "Cryo-OrbiSIMS for 3D molecular imaging of a bacterial biofilm in its native state"

Junting Zhang, et al.

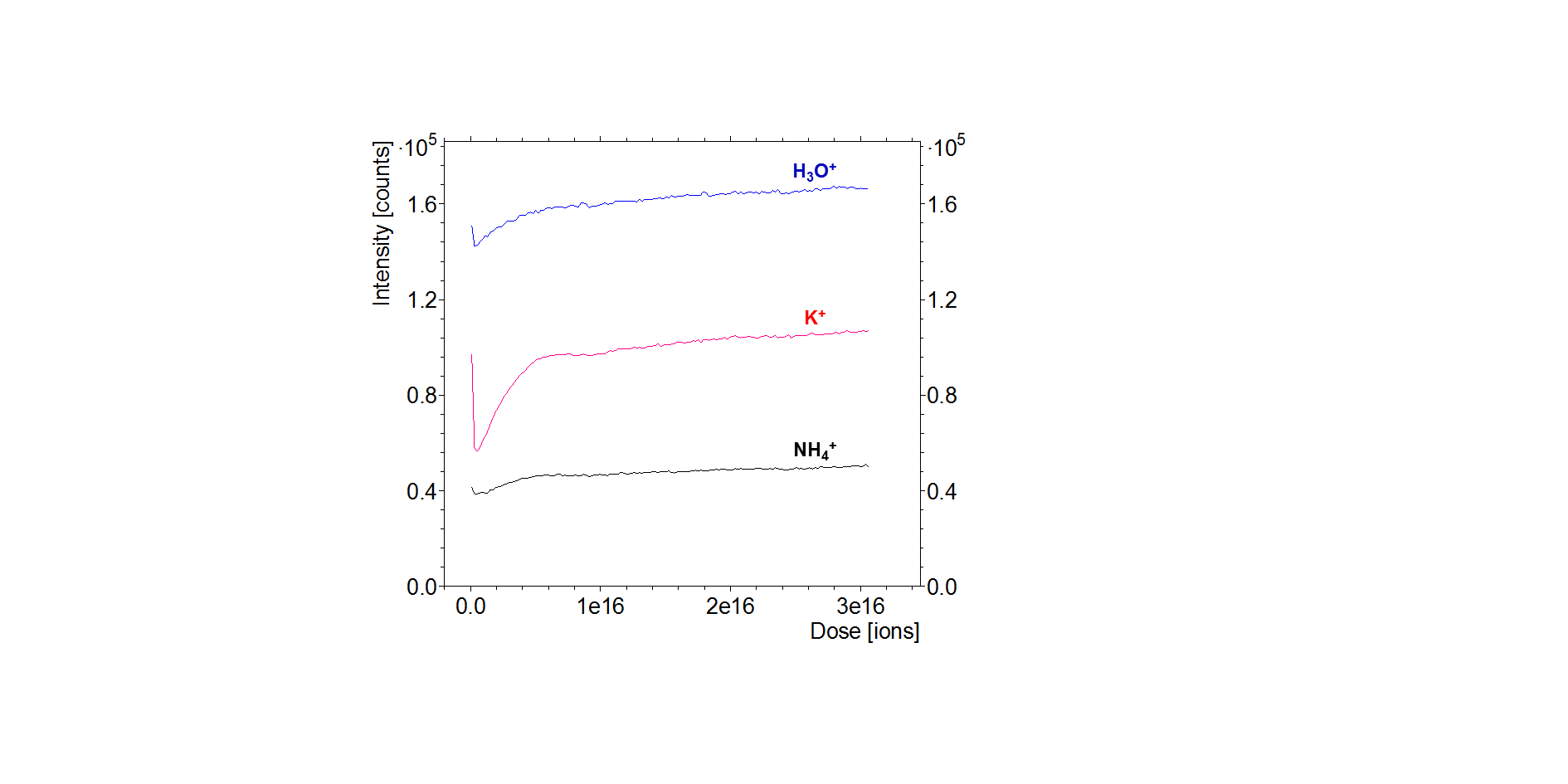

**Supplementary Figure 1**. 30 keV Bi_3_^+^ ToF MS positive ion intensity depth profile (mode 10) of frozen-hydrated *P. aeruginosa* biofilm for H_3_O^+^ at *m/z* 19 (blue line), K^+^ at *m/z* 39 (red line), NH_4_^+^ at *m/z* 18 (grey line).

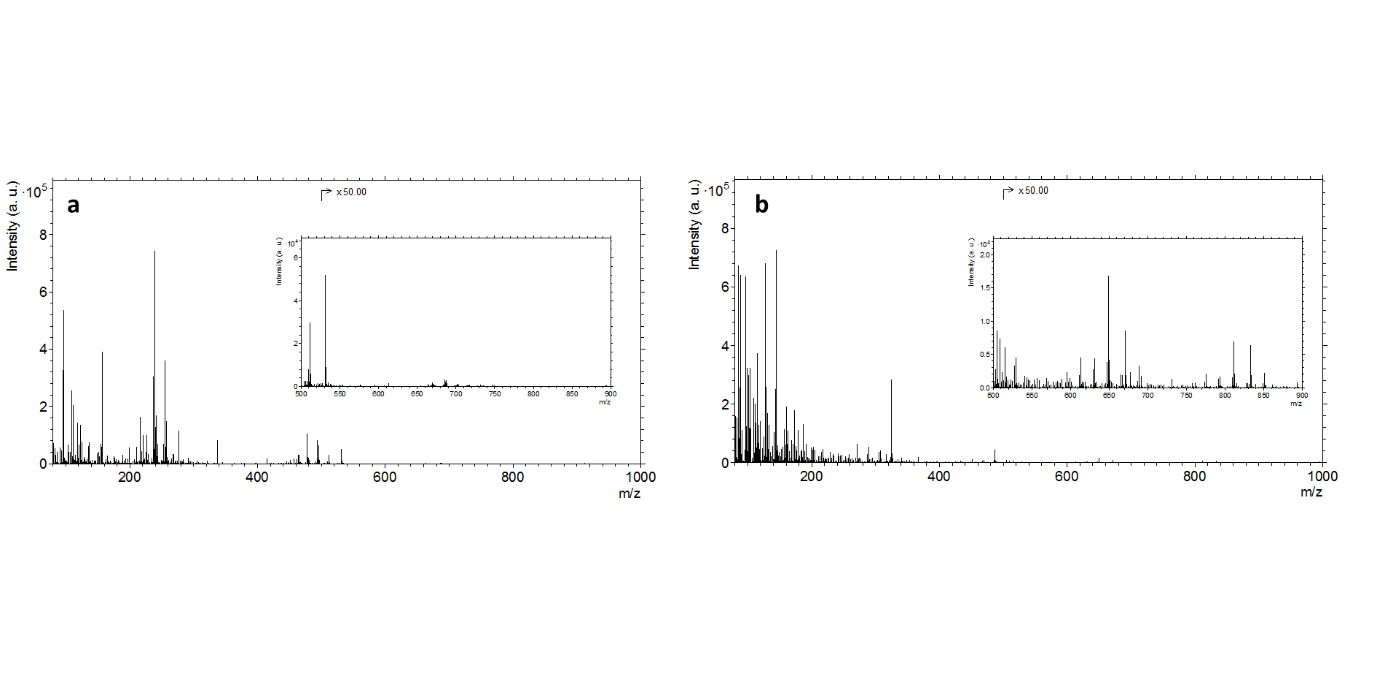

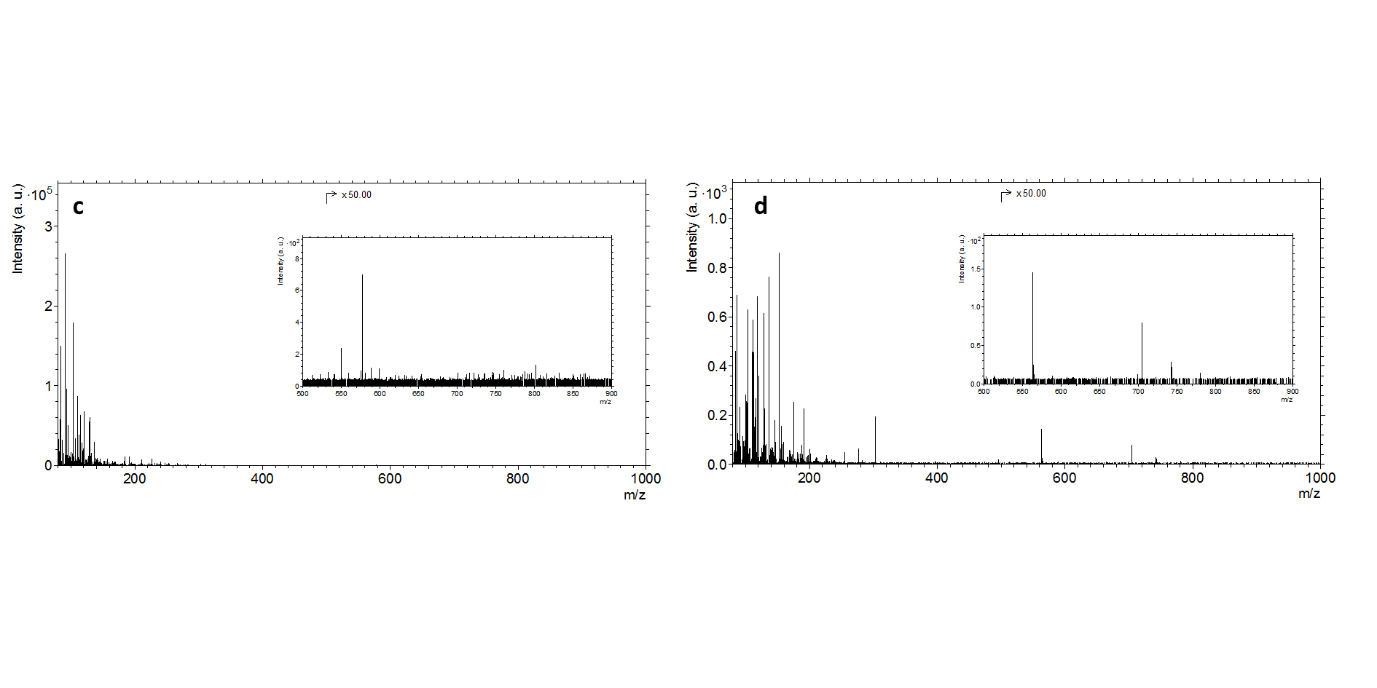

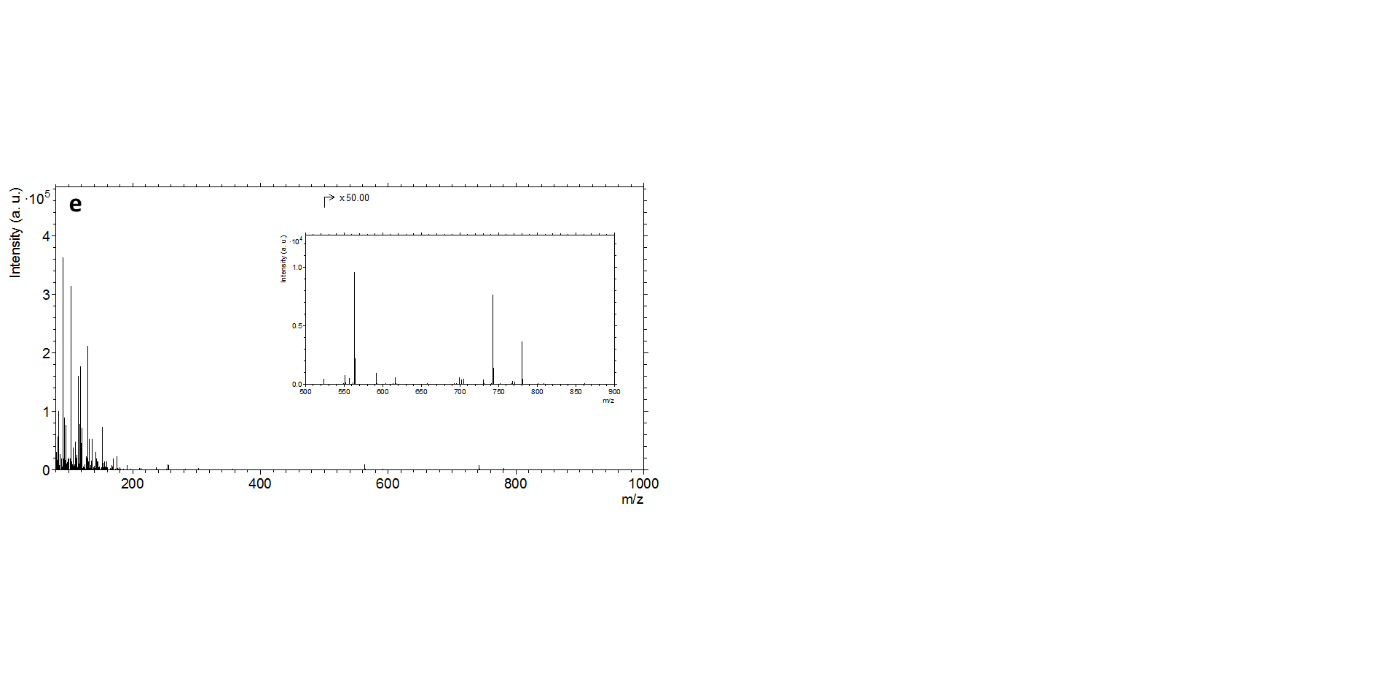

**Supplementary Figure 2.** 20 keV Ar_3000_^+^ GCIB Orbitrap positive ion MS (mode 4) of a frozen-hydrated *E. coli* cell pellet with different cryo-protectants. (a) 1-hexadecene, (b) 20% dextran, (c) 5% BSA, (d) 20% methanol**,** (e) 150 mM ammonium formate. Insets shows details from *m/z* 500-900 region.

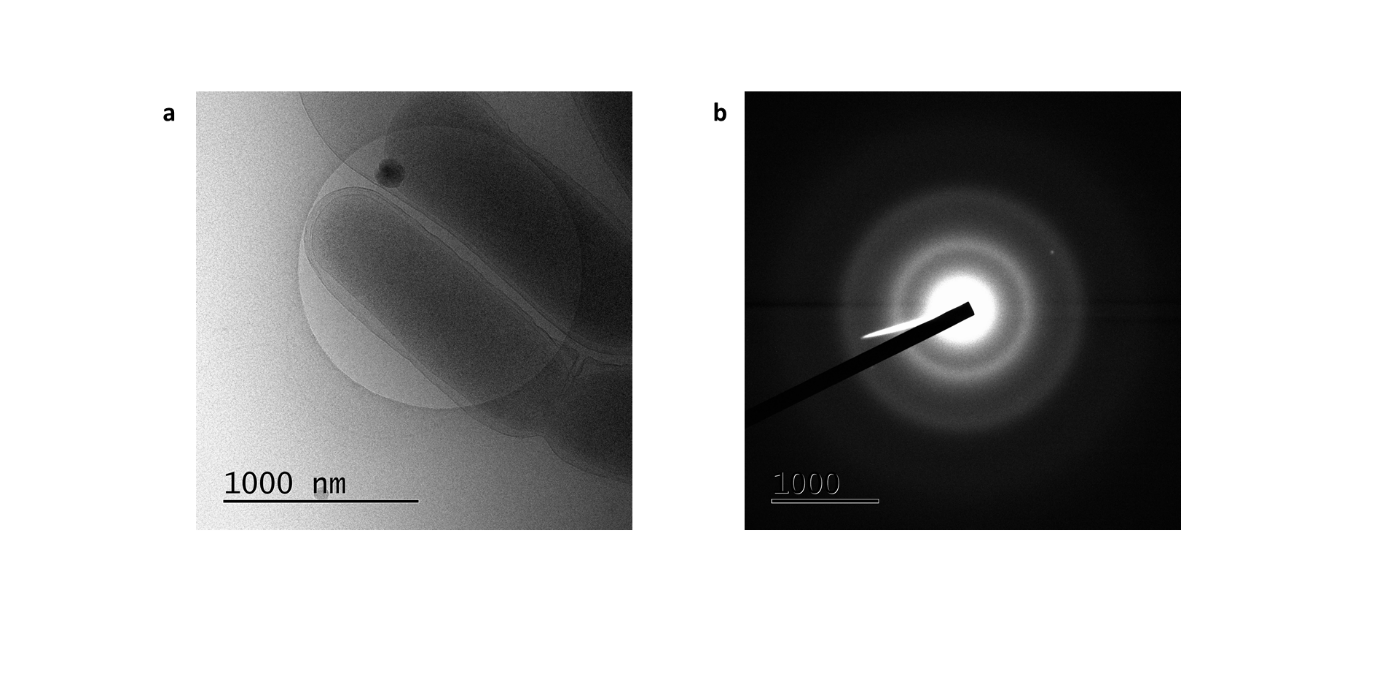

**Supplementary Figure 3.** Cryo transmission electron micrography (a) of planktonic *P. aeruginosa* in 150mM ammonium formate, taken at 20,000 x magnification and (b) electron diffraction pattern of the same area.

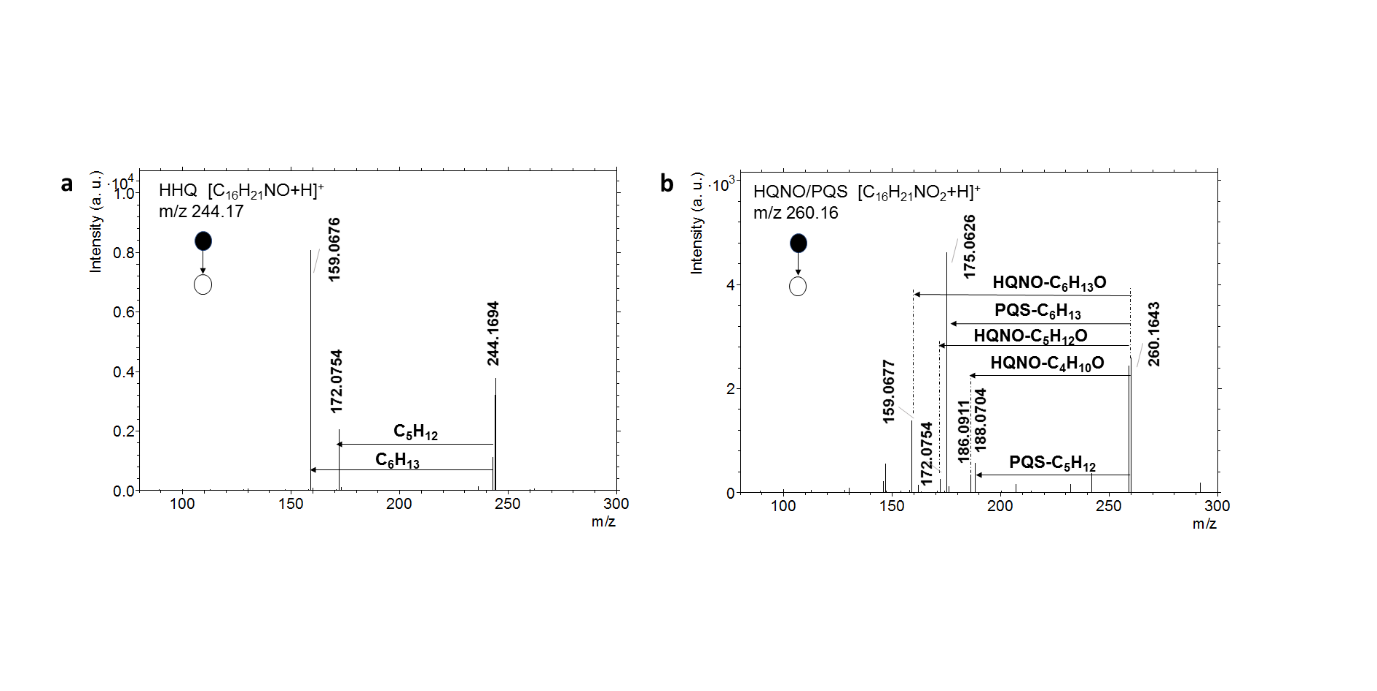

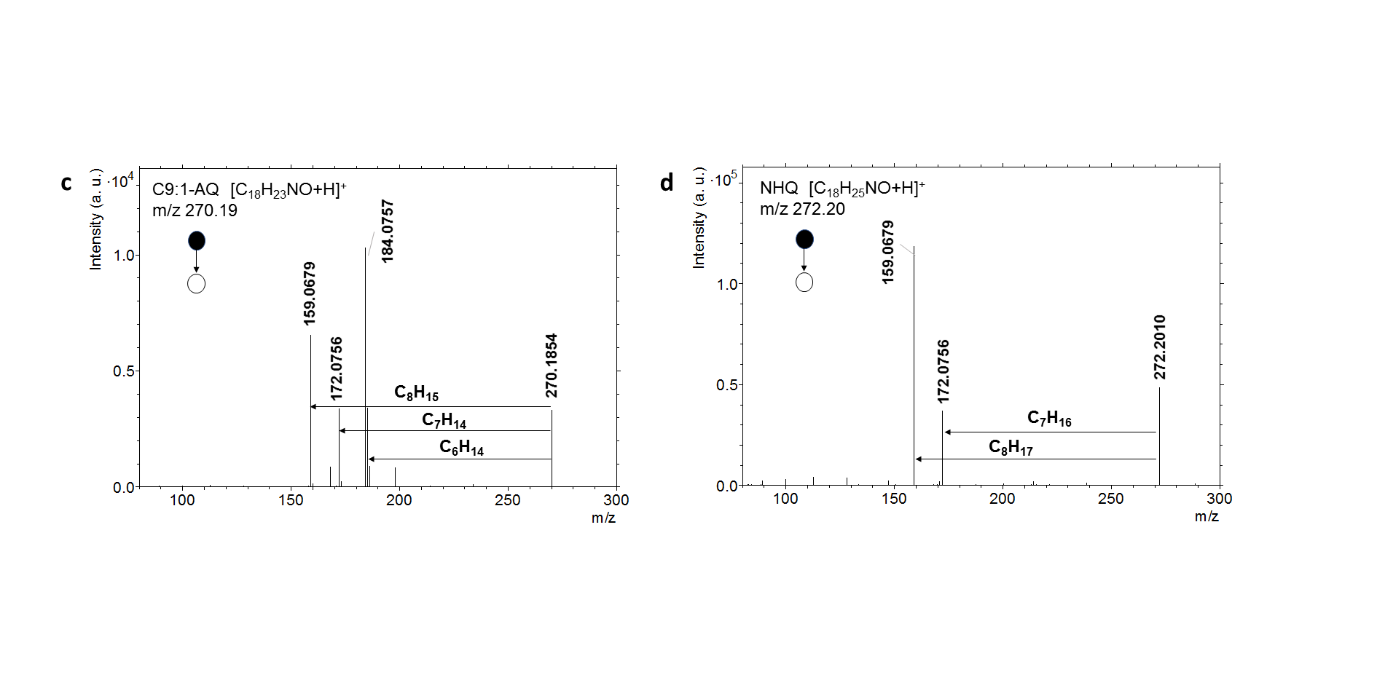

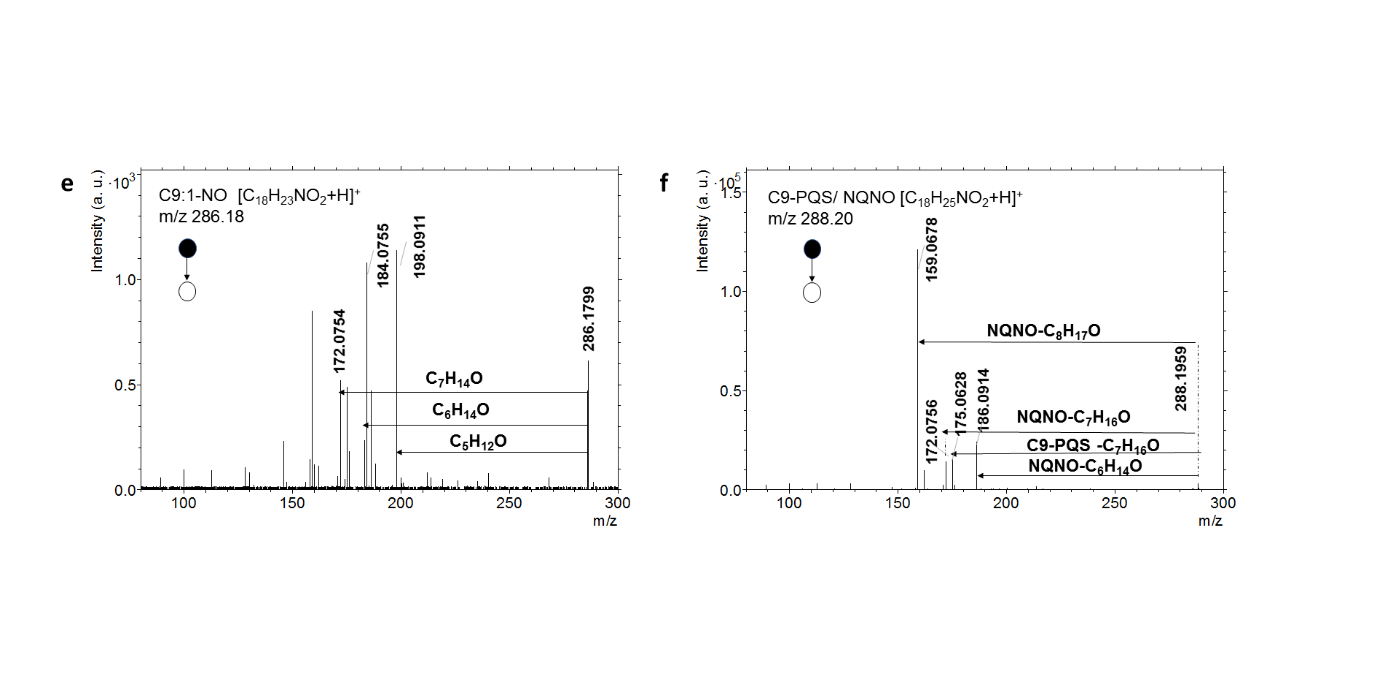

**Supplementary Figure 4.** 20 keV Ar_3000_^+^ GCIB Orbitrap positive MS/MS (mode 2) of frozen hydrated *P. aeruginosa* biofilm. (a) HHQ [M+H]^+^, (b) HQNO/C7-PQS [M+H]^+^, (c) C9:1-AQ [M+H]^+^, (d) NHQ [M+H]^+^, (e) C9:1-NO and (f) C9-PQS/NQNO

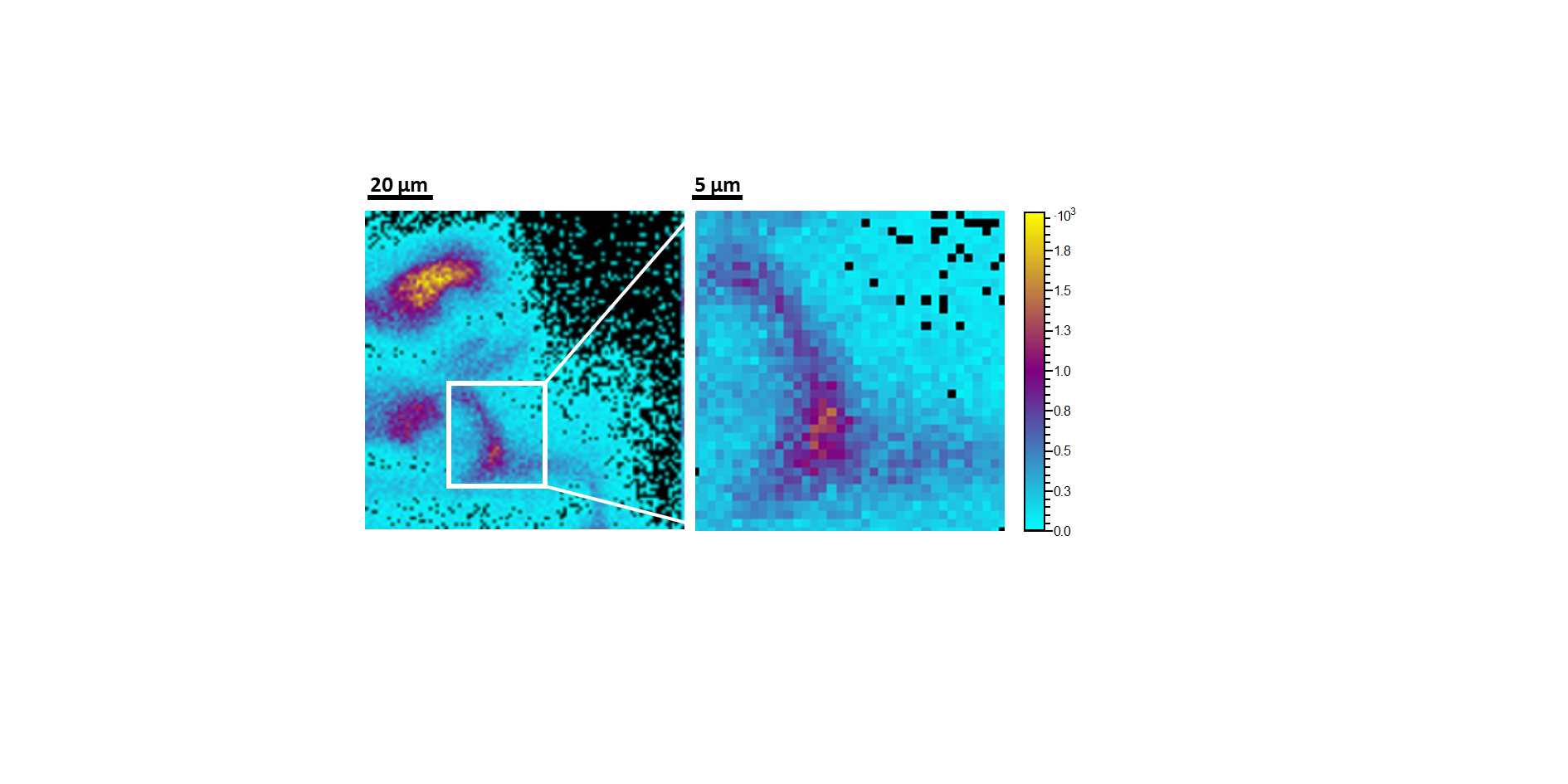

**Supplementary Figure 5.** 20 keV Ar_3000_^+^ 2D Orbitrap MS positive ion image (mode 10), acquired with 1 µm pixel size, for adenine (m/z 136.0617, C_5_H_6_N_5_^+^) in frozen-hydrated *P. aeruginosa* biofilm.

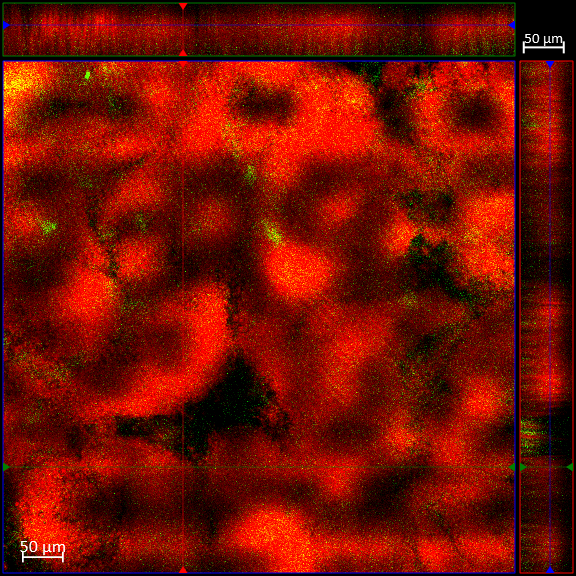

**Supplementary Figure 6.** Confocal laser scanning microscopy (CLSM) of a *Pseudomonas aeruginosa* biofilm grown on aluminium sample carriers. The biofilms were visualised via constitutive expression in *P. aeruginosa* cells of the red fluorescent protein mCherry and staining of the extracellular DNA (eDNA) with the dye YOYO-1 (green). The central image shows a top-down view of the biofilm; side panels are vertical sections

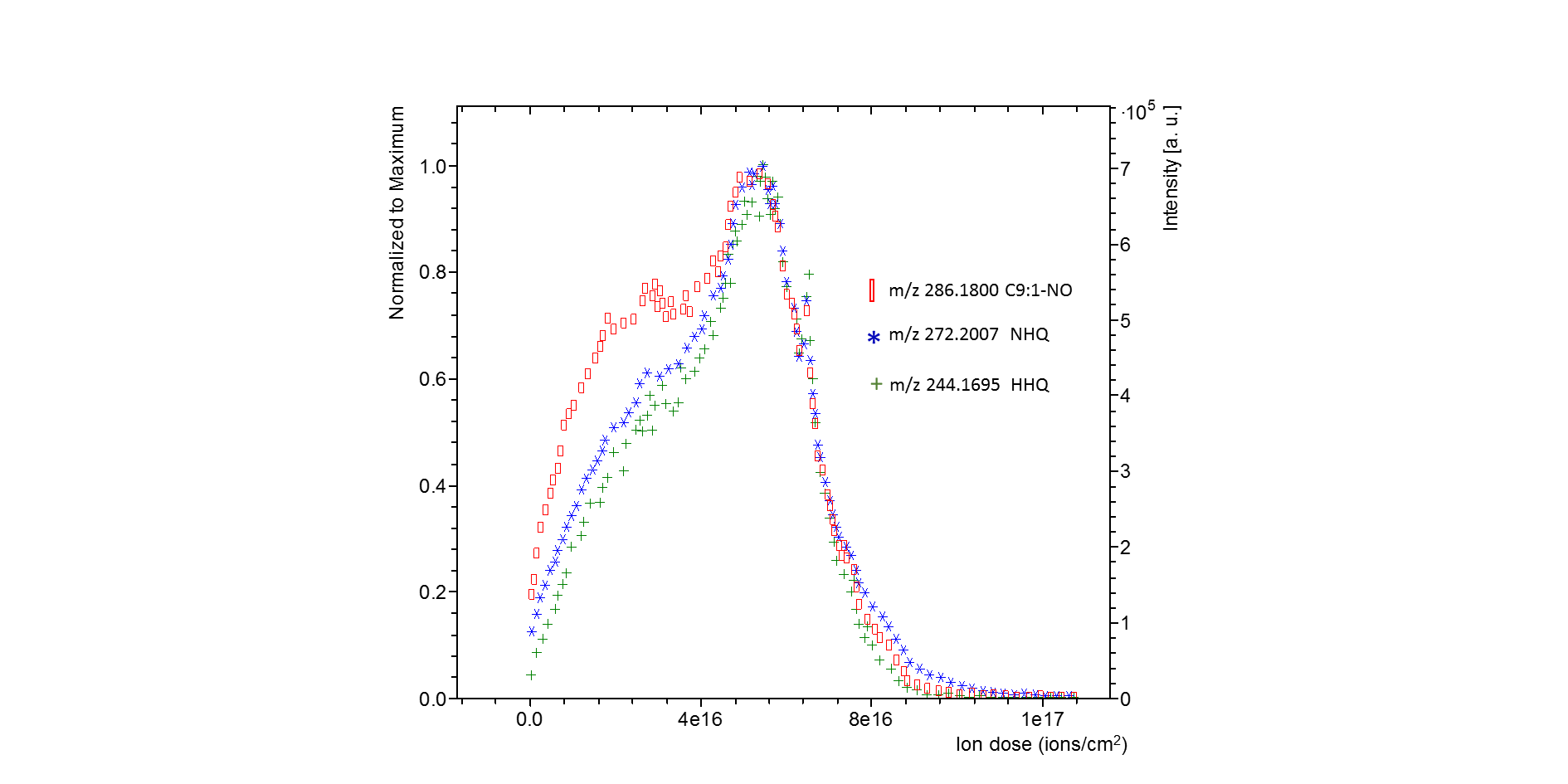
**Supplementary Figure 7.** 20 keV Ar_3000_^+^ GCIB Orbitrap MS positive ion intensity depth profile (mode 4) of frozen-hydrated biofilm for 3 alkyl quinolone signals: HHQ (C_16_H_22_NO, *m/z* 244.1695), NHQ (C_18_H_26_NO, *m/z* 272.2007) and C9:1-NO (C_18_H_24_NO_2_, m/z 286.1800). All the signals are normalized to the maximum and summed over a 300 × 300 µm area.

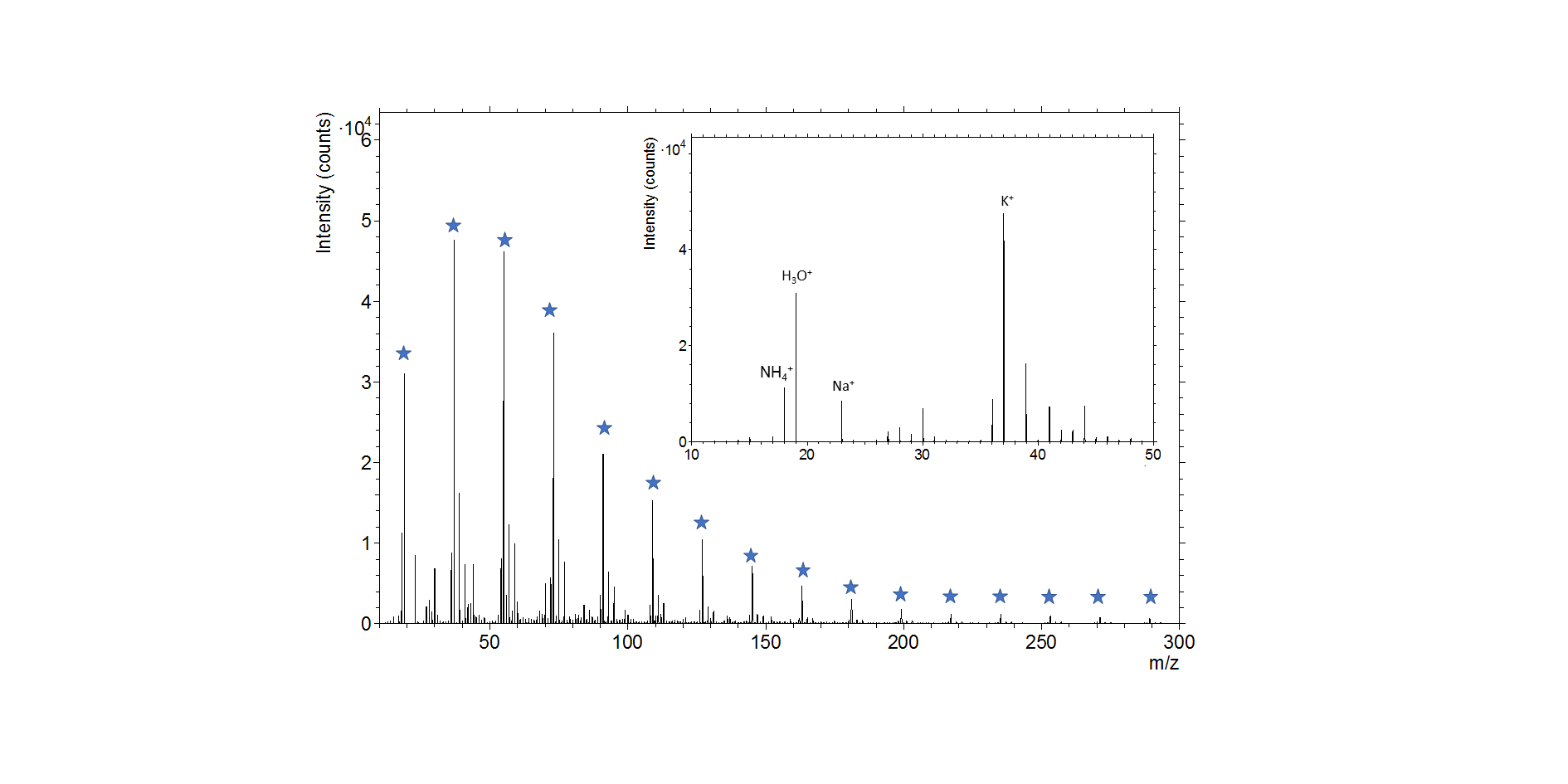

**Supplementary Figure 8.** 30 keV Bi_3_^+^ ToF MS positive ion spectrum (mode 10) of frozen-hydrated *P. aeruginosa* biofilm. Star symbols indicate water cluster ions, with the formula [H(H_2_O)_n_]^+^. Detail is shown in the inset (*m/z* 10 - *m/z* 50). The spectrum is the sum of 150 scans with a total ion dose of 1.36 × 10^16^ ions/cm^2^.

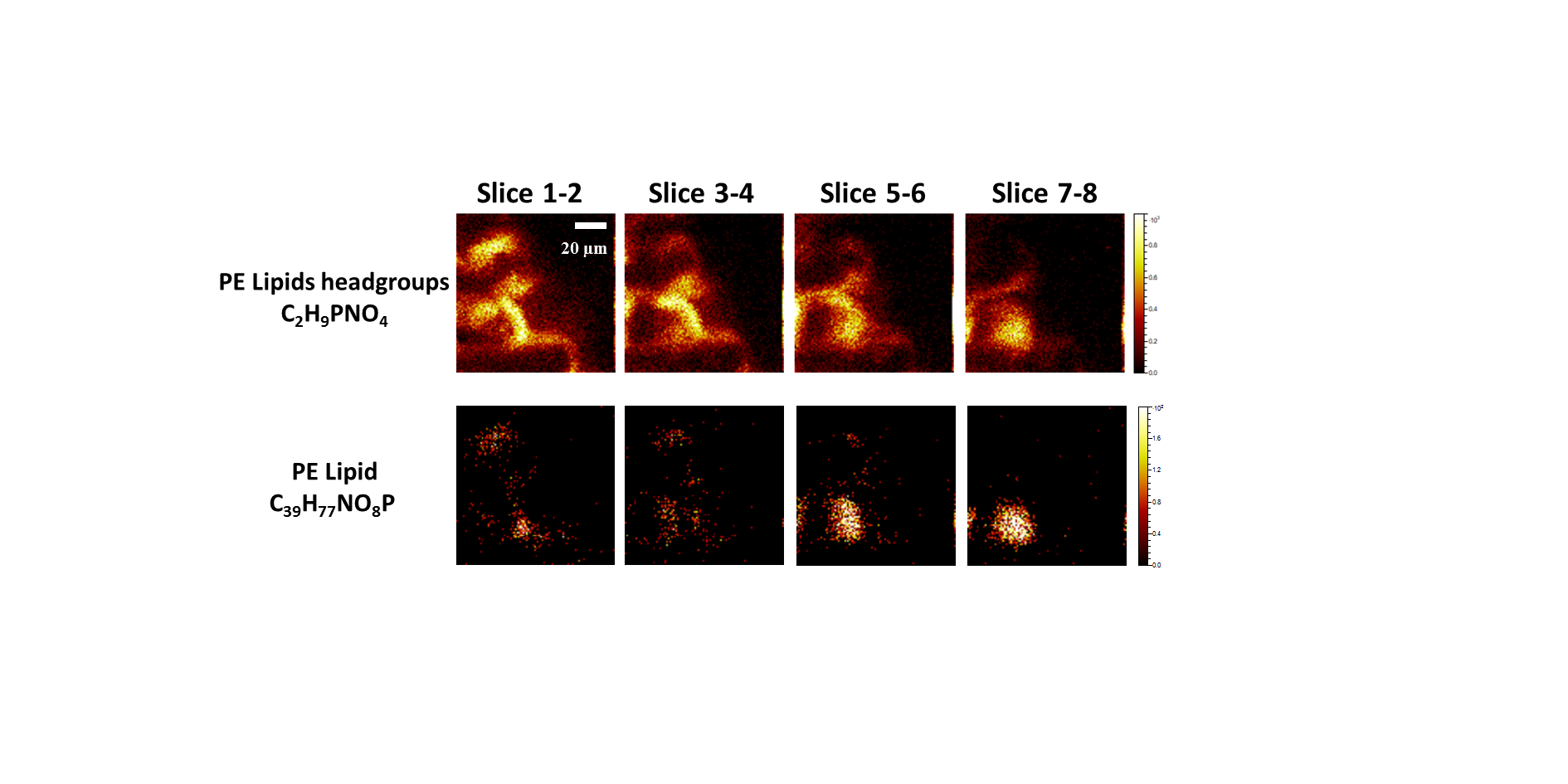

**Supplementary Figure 9.** 3D Orbitrap MS images (mode 8) of frozen-hydrated *P. aeruginosa* biofilm. Sequence of 20 keV Ar_3000_^+^ (pixel size 1 µm) Orbitrap MS images of PE lipid headgroups (C_2_H_9_PNO_4_^+^ at *m/z* 142.0262) and PE lipid (C_39_H_77_NO_8_P^+^ at *m/z* 718.5377)

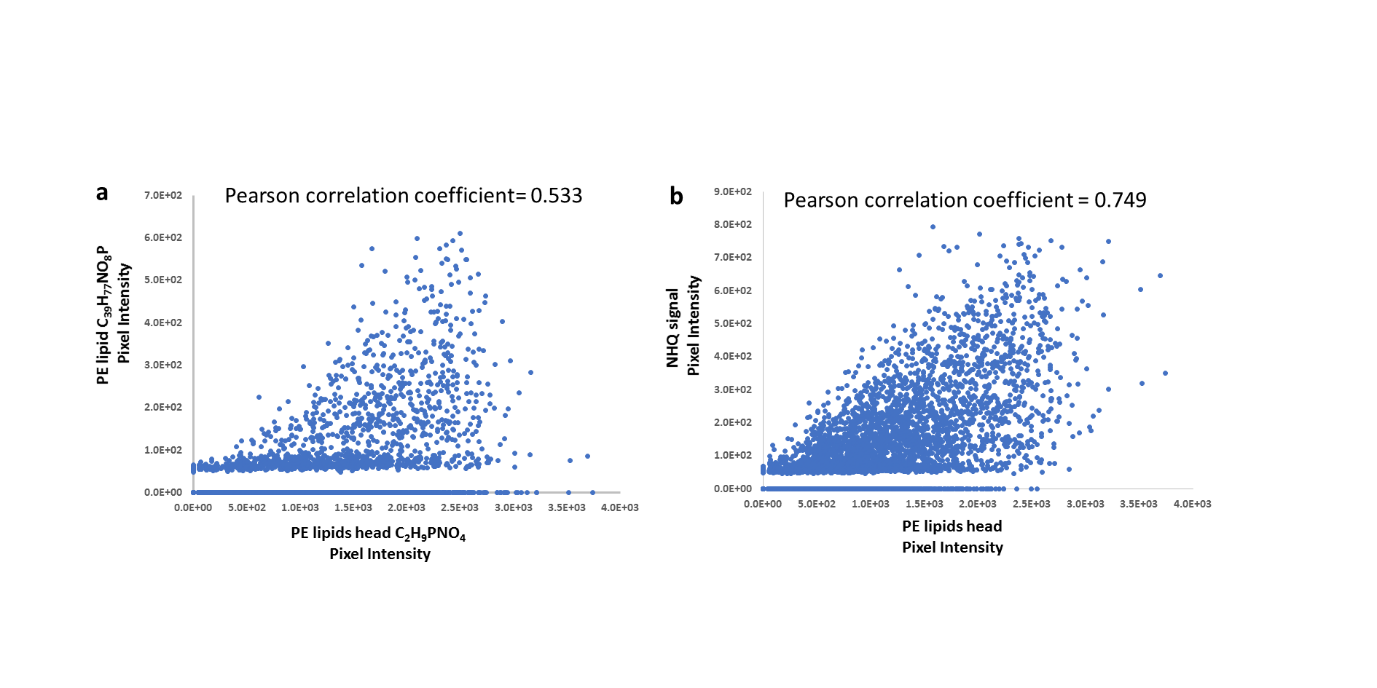

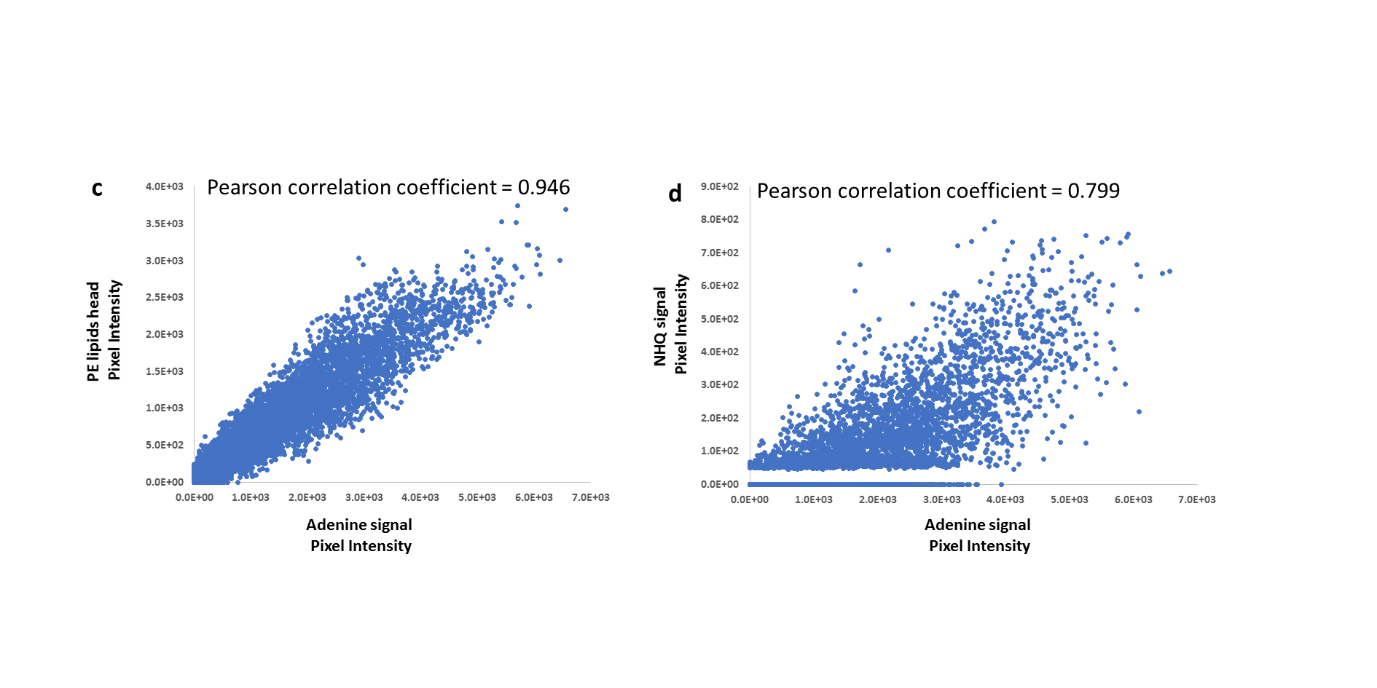

**Supplementary Figure 10.** Scatter plot of pixel intensity of (a) PE lipid (C_39_H_77_NO_8_P^+^, at *m/z* 718.5380) to PE lipids head (C_2_H_9_PNO_4_^+^, at *m/z* 142.0262); (b) uracil (C_4_H_5_N_2_O_2_^+^ at *m/z* 113.0344) to adenine (C_5_H_6_N_5_^+^ at *m/z* 136.0617); (c) PE lipids head (C_2_H_9_PNO_4_^+^, at *m/z* 142.0262) to adenine (C_5_H_6_N_5_^+^ at *m/z* 136.0617); (d) NHQ (C_18_H_26_NO^+^ at *m/z* 272.2007) to adenine (C_5_H_6_N_5_^+^ at *m/z* 136.0617); (e) NHQ (C_18_H_26_NO^+^ at *m/z* 272.2007) to PE lipids head (C_2_H_9_PNO_4_^+^, at *m/z* 142.0262). The data exported from Orbitrap images (same data used in **Supplementary Fig. 9)** which sums 9 scans together.

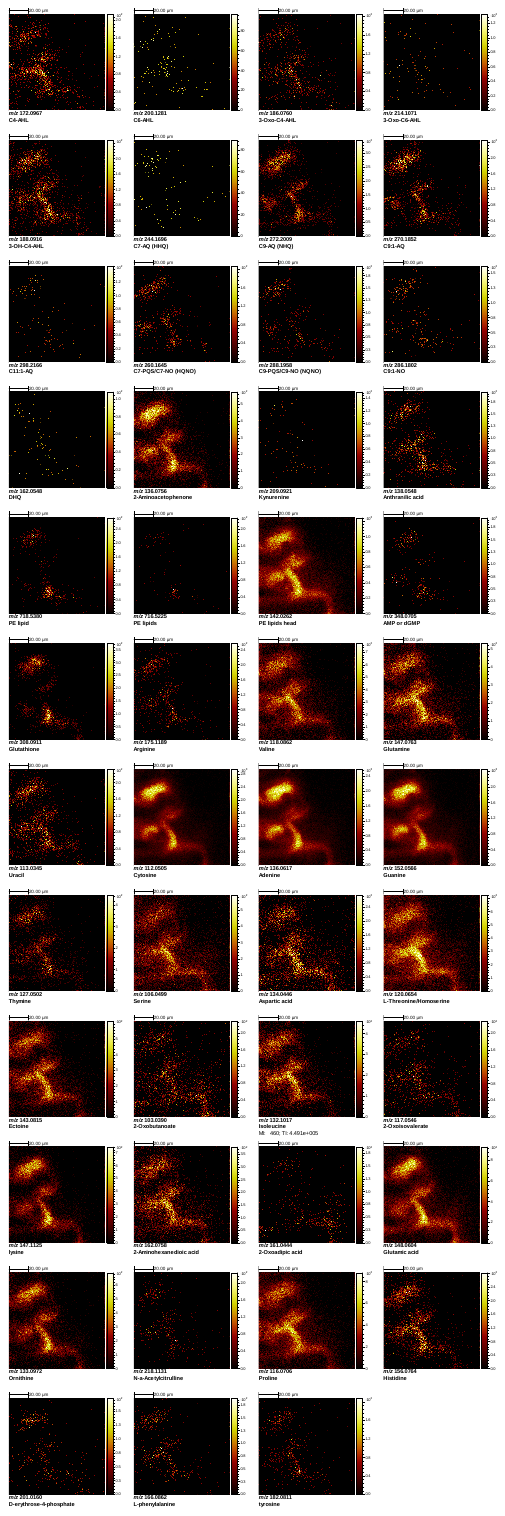

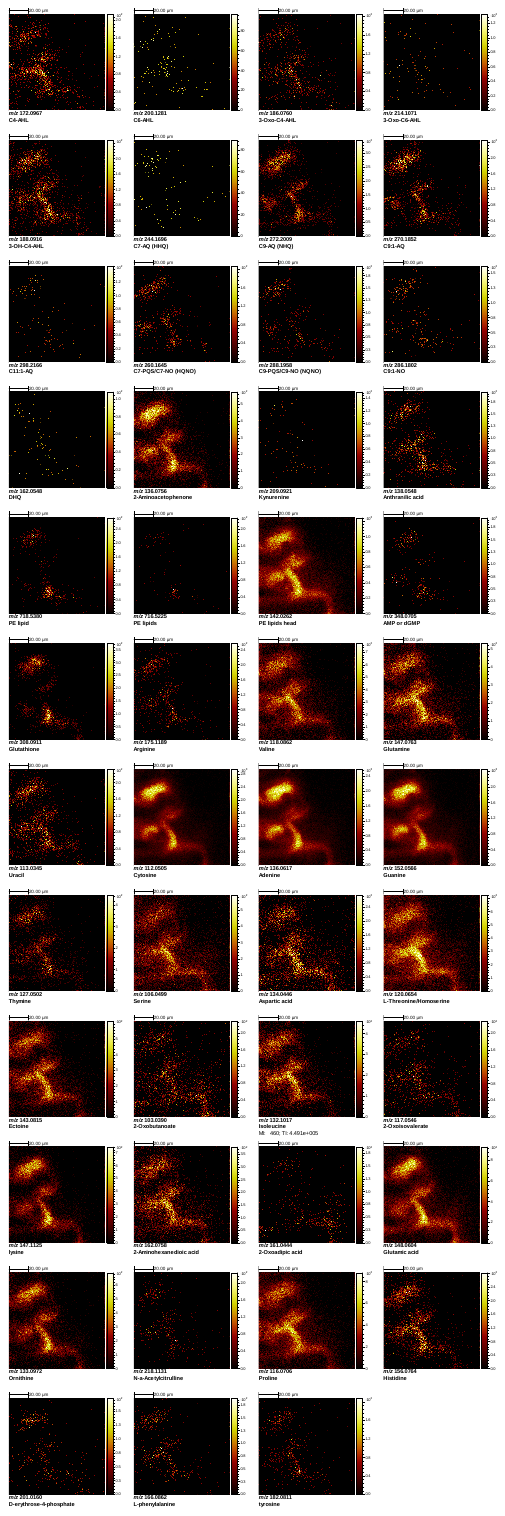

**Supplementary Figure 11.** 20 keV Ar_3000_^+^ (pixel size 1µm) Orbitrap MS images (mode 8) showing the distribution of different compounds in the 100 × 100 µm area of frozen-hydrated *P. aeruginosa* biofilm. The scale bar is 20 µm.

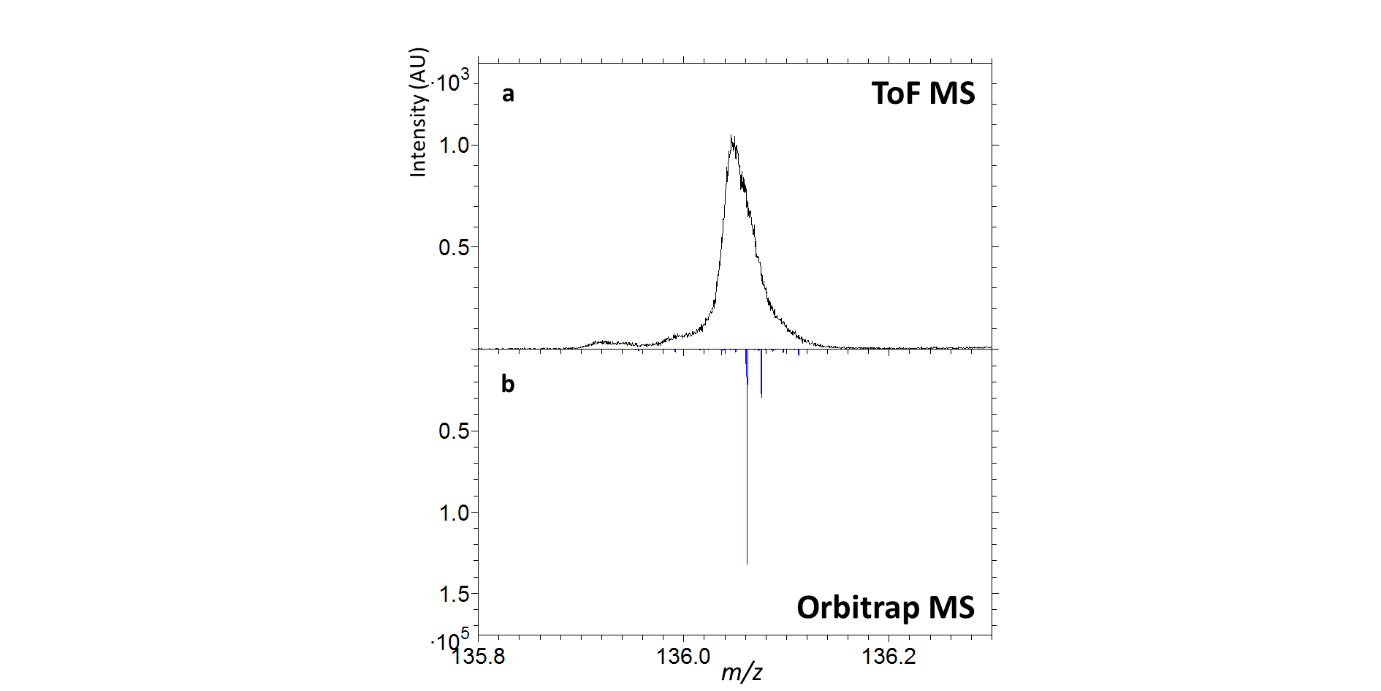

**Supplementary Figure 12.** A comparison between 20 keV Ar_3000_^+^ GCIB Orbitrap MS positive ion spectrum and 30 keV Bi_3_^+^ ToF MS positive ion spectrum (*m/z* 135 - *m/z* 137) (mode 10) of frozen-hydrated *P. aeruginosa* biofilm. (a) Low-mass-resolution spectrum obtained during imaging with the ToF MS and (b) high-mass-resolution spectrum from sputtered material using Orbitrap MS which is composed of two peaks: adenine (m/z 136.0617) and 2-aminoacetophenone (*m*/*z* 136.0756).

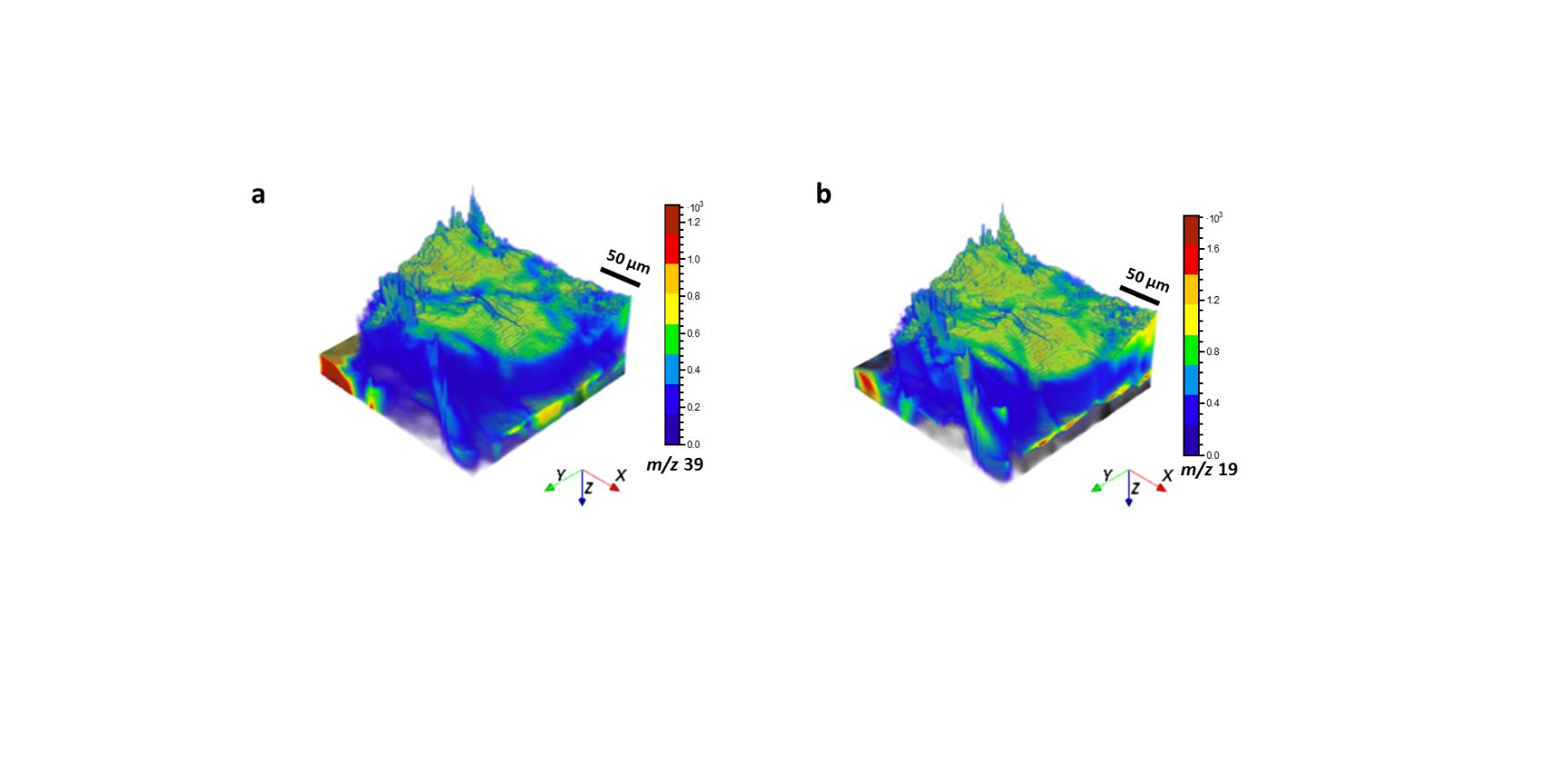

**Supplementary Figure 13.** 3D rendered ToF MS image (mode 10) of frozen-hydrated *P. aeruginosa* biofilm. (a) K^+^ (*m/z* 39) on the substrate (Al, m*/z* 27, shown in grey); (b) H_3_O^+^ (*m/z* 19) on the substrate (Al, m*/z* 27, shown in grey); The image was corrected assuming the substrate is flat.

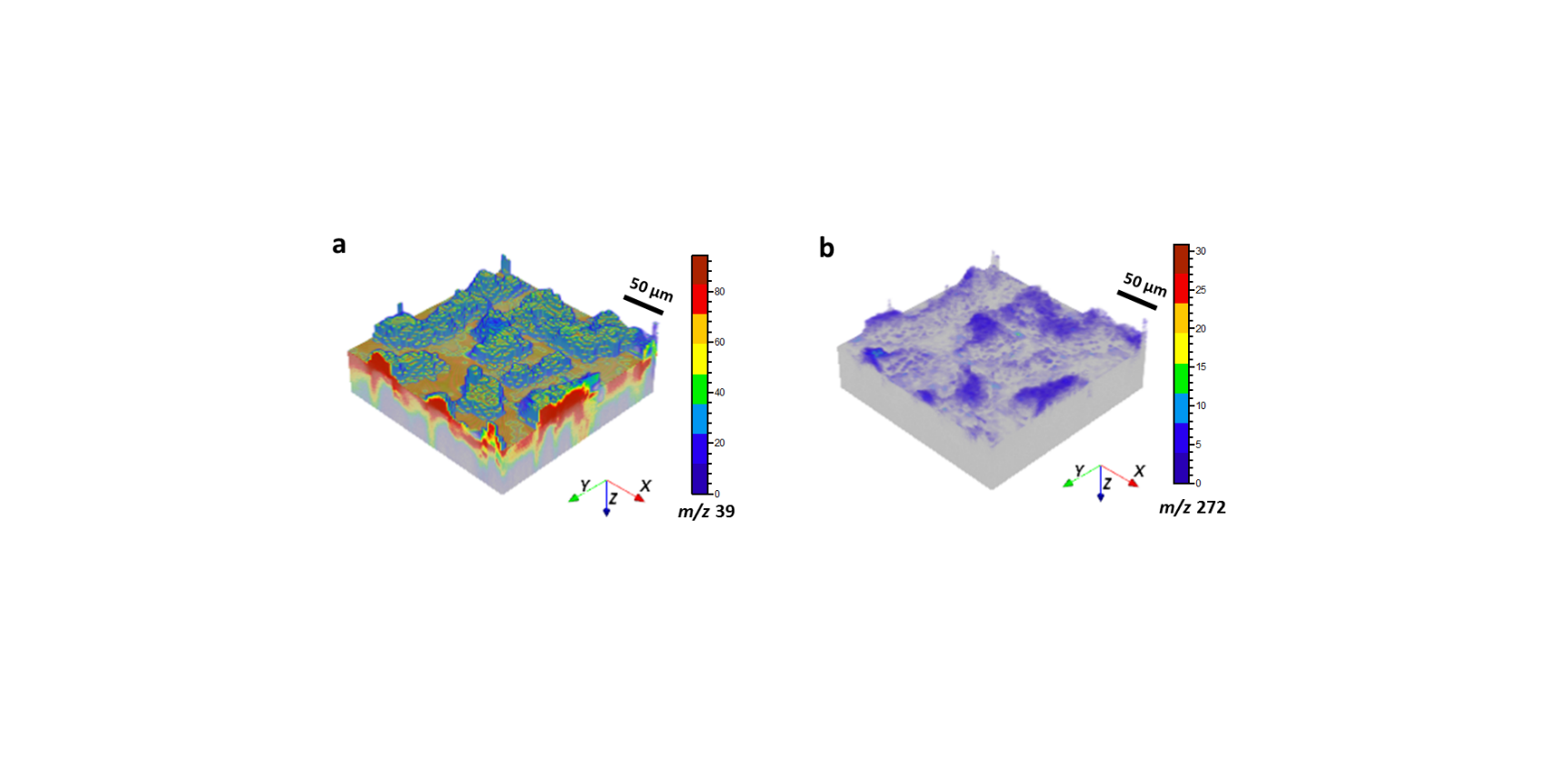

**Supplementary Figure 14.** 3D rendered ToF MS image (mode 10) of freeze-dried *P. aeruginosa* biofilm. (a) K^+^ (*m/z* 39) on the substrate (Al, m*/z* 27, shown in grey); (b) NHQ (*m/z* 272) on the substrate (Al, *m/z* 27). The image was corrected assuming the substrate is flat.

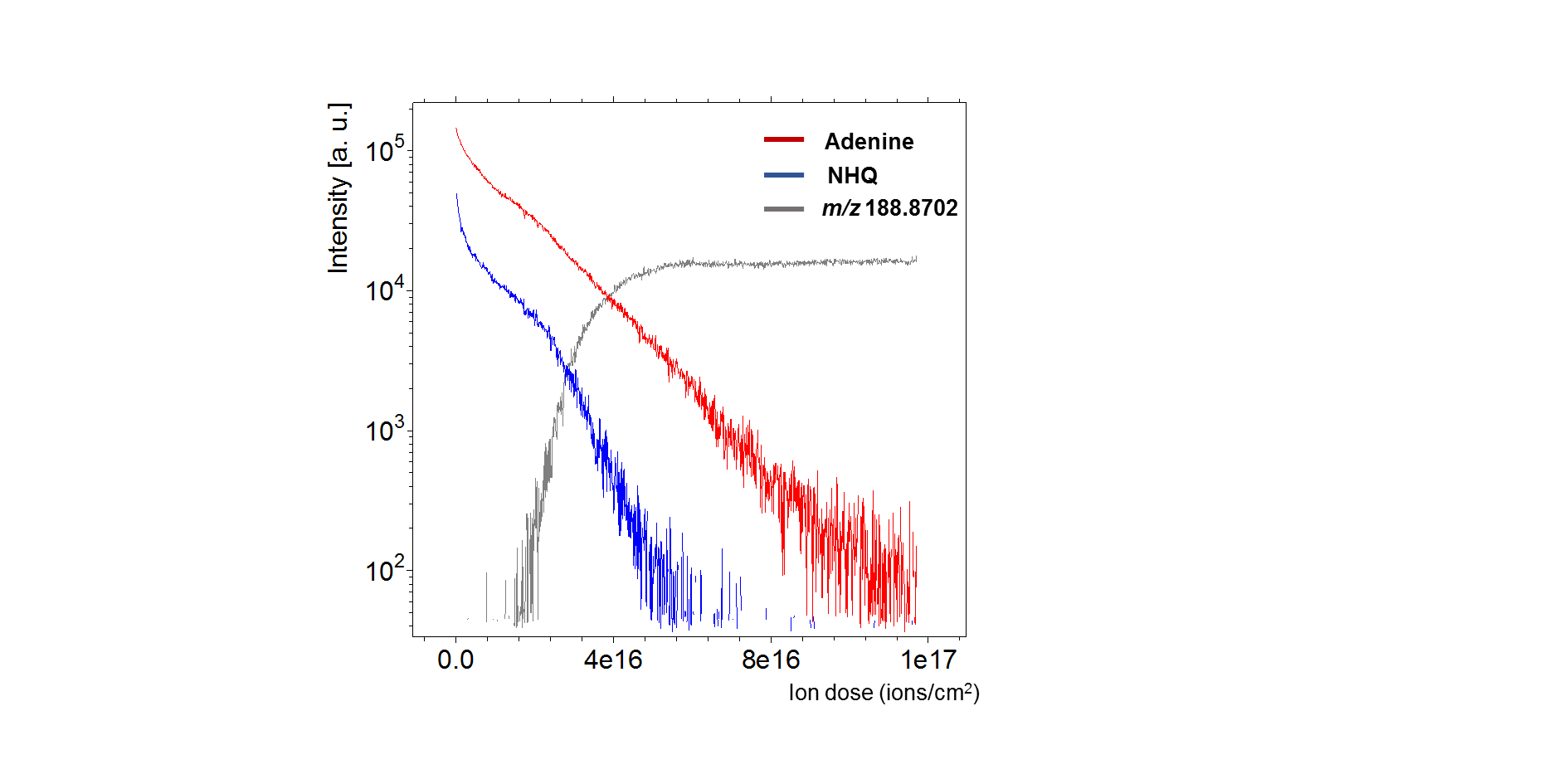

**Supplementary Figure 15.** 20 keV Ar _3000_^+^ GCIB Orbitrap MS positive ion intensity depth profile (mode 4) of freeze-dried *P. aeruginosa* biofilm with adenine at *m/z* 136.0618 (red line), NHQ at *m/z* 272.2007 (blue line), substrate-related signal at *m/z* 188.8702.

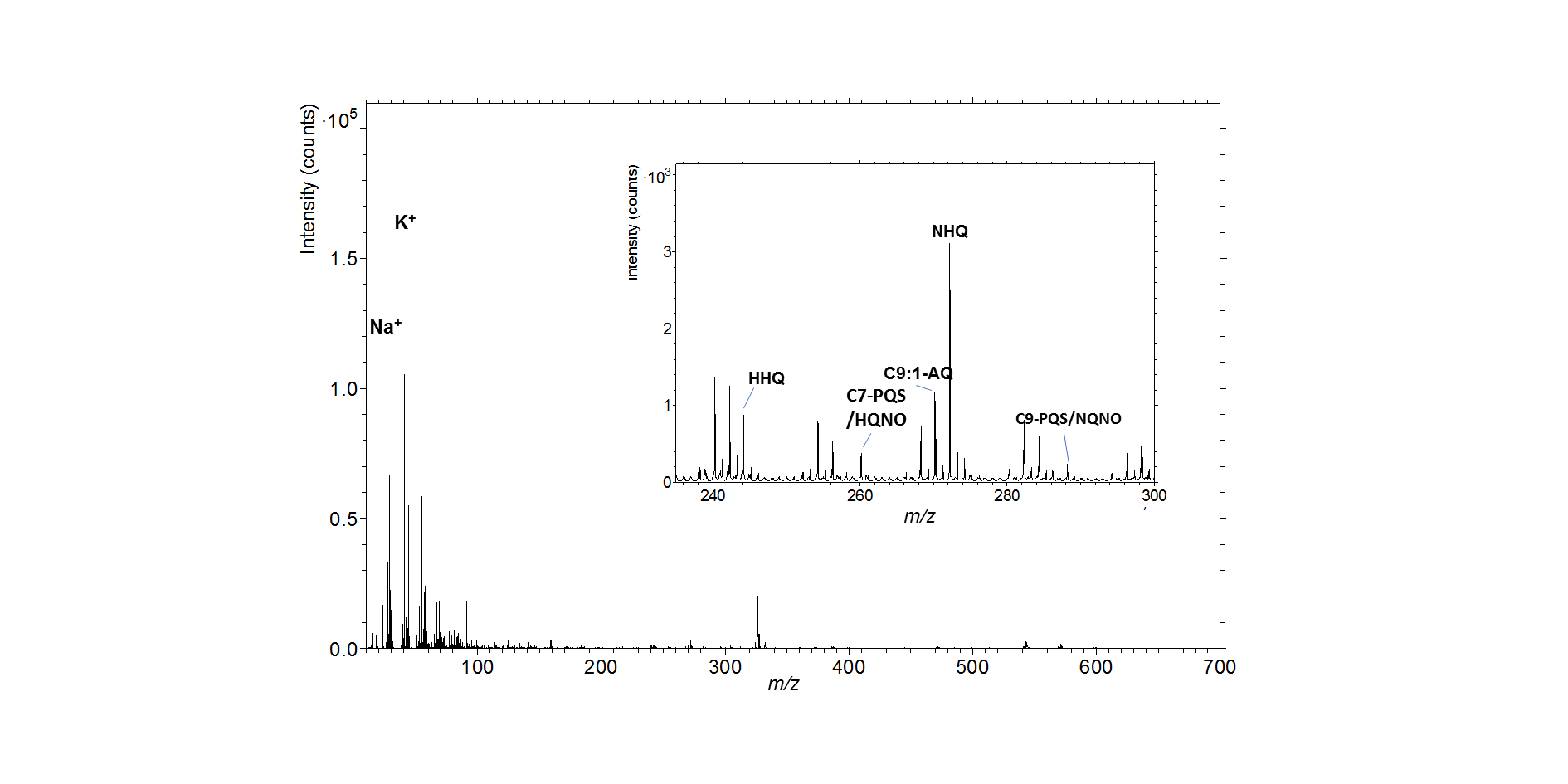

**Supplementary Figure 16**. 30 keV Bi_3_^+^ ToF positive ion MS (mode 10) of freeze-dried *P. aeruginosa* biofilm. In the inset (*m/z* 240 - *m/z* 300), some alkyl quinolones related signals are putatively annotated.

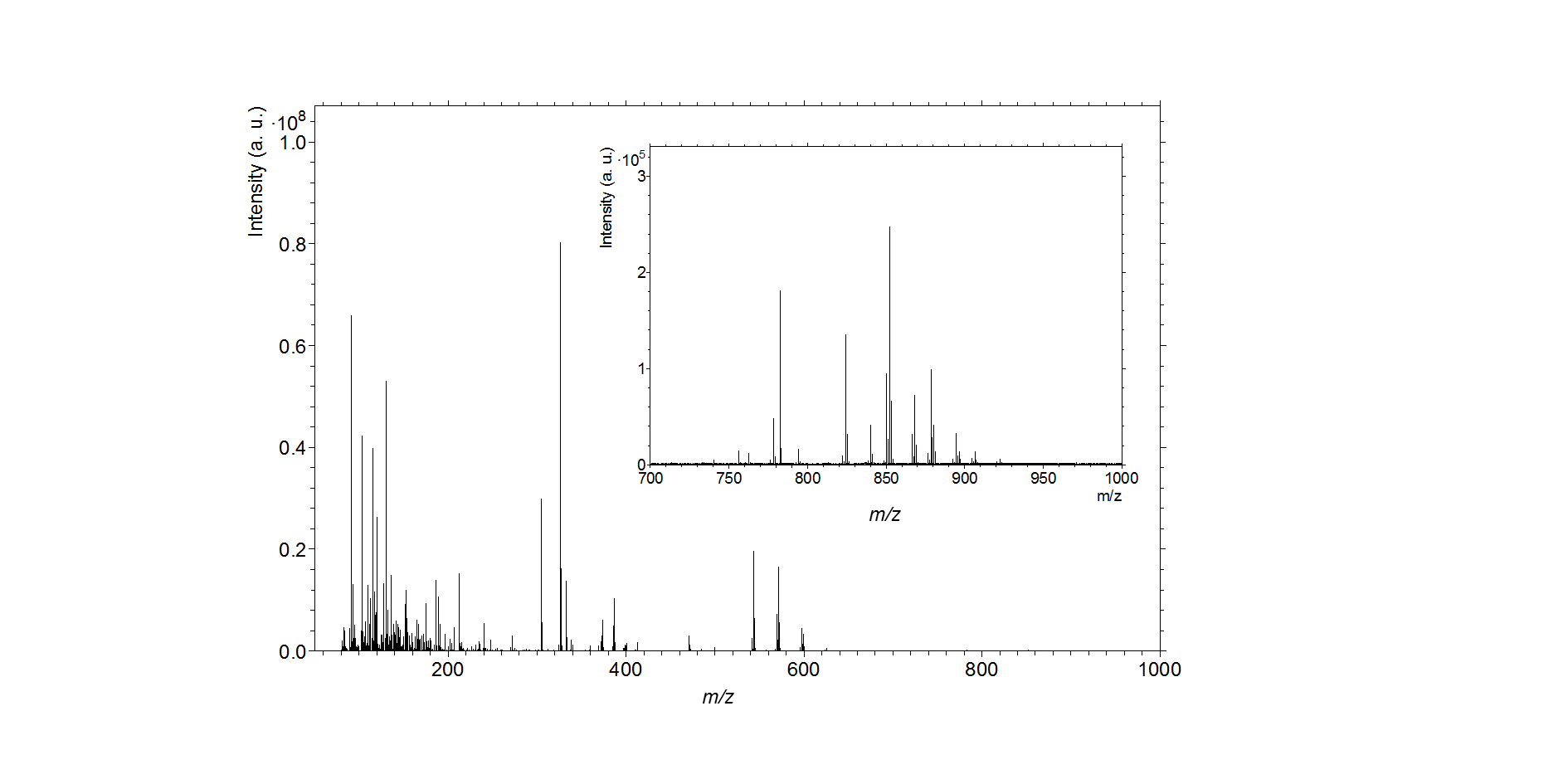

**Supplementary Figure 17.** 20keV Ar_3000_ ^+^ GCIB Orbitrap positive ion MS (mode 4) of freeze-dried *P. aeruginosa* biofilm. Inset (m/z 700-1000) shows low intensity ions of PE lipids and other ions in the m/z 800-900 that are not annotated.

frozen enhancement ratio

| Xlog P | amino acids |
| --- | --- |
| -3.1 | 27673.4 |
| -5.6 | 1299.36 |
| -2.9 | 48246.2 |
| -2.5 | 55.0091 |
| -3.4 | 50.2871 |
| -1.9 | 312.26 |
| -1.7 | 20321.9 |
| -3 | 95392.1 |
| -3.7 | 73401.8 |
| -4.4 | 53148.8 |
| -2.3 | 3279.63 |
| -2.5 | 11842.1 |
| -3.2 | 1855.92 |
| -4.2 | 4065.29 |
| -1.1 | 231.244 |
| -1.5 | 527.745 |
| -2.3 | 3073.51 |
| -3.1 | 77.6613 |
| -2.3 | 1859.79 |
| -3.1 | 27076.3 |

| Xlog P | nucleobases |
| --- | --- |
| -0.6 | 582.252 |
| -1.1 | 436.445 |
| -1.7 | 48.2096 |
| -0.1 | 16.022 |
| -1.1 | 42.6974 |

| Xlog P | quinolones |
| --- | --- |
| 3.3 | 18.4463 |
| 4.59 | 8.85114 |
| 4.9 | 9.59112 |
| 5.42 | 6.39318 |
| 6 | 11.9755 |
| 6.26 | 1.38161 |
| 6.5 | 17.7559 |
| 4.5 | 60.1911 |
| 4.08 | 59.4129 |
| 4.63 | 33.4498 |
| 5.45 | 73.395 |
| 2.72 | 82.4716 |
| 3.17 | 40.3109 |
| 5 | 29.1742 |
| 3.18 | 616.571 |
| 4.1 | 66.3571 |
| 5.01 | 59.2526 |
| 2.59 | 362.483 |
| 3.43 | 303.016 |
| 2.41 | 34.4591 |
| 3.25 | 135.229 |

| Xlog P | lactones |
| --- | --- |
| 0.5 | 4836.59 |
| 1.5 | 1076.82 |
| 0.9 | 396.08 |
| -0.1 | 16998.6 |

| Xlog P | other metabolites |
| --- | --- |
| 1.6 | 2.77405 |
| 0.8 | 12.1196 |
| -2.2 | 363.663 |
| 1.2 | 20.9809 |
| -2.6 | 1502.98 |
| -3.5 | 1164.67 |
| -0.8 | 2906.13 |
| -0.6 | 9176.63 |
| 0.1 | 12032.3 |
| 0.7 | 5395.44 |
| -3.1 | 23733.9 |
| -1.1 | 145.812 |
| -3.3 | 5748.91 |
| -4.5 | 22296.2 |
| -0.5 | 4508.52 |

**Supplementary Figure 18.** Scatter plot of the positive signal enhancement ratio (frozen hydrated / freeze dried) with the XLogP values for amino acids, nucleobases, quinolones, lactones and other metabolites from the biofilm. Data is from Supplementary Table 2. Lipids are not plotted. The vertical axis is a log scale.

**Supplementary Table 1.** Putative annotation of compounds in the positive ion spectrum of frozen-hydrated *P. aeruginosa* biofilm.

| **Compounds** | **Adduct** | **Chemical Formula** | ***m*/*z*** | **Mass Accuracy (ppm)** |
| --- | --- | --- | --- | --- |
| **2-alkyl-4-quinolones** | | | | |
| C5-AQ | [M+H]^+^ | C_14_H_18_NO | 216.1379 | -1.8 |
| C6-AQ | [M+H]^+^ | C_15_H_20_NO | 230.1540 | 0.3 |
| C7-AQ (HHQ) | [M+H]^+^ | C_16_H_22_NO | 244.1696 | 0.0 |
| C8-AQ | [M+H]^+^ | C_17_H_24_NO | 258.1851 | -0.7 |
| C9-AQ (NHQ) | [M+H]^+^ | C_18_H_26_NO | 272.2009 | -0.1 |
| C10-AQ | [M+H]^+^ | C_19_H_28_NO | 286.2174 | 3.0 |
| C11-AQ | [M+H]^+^ | C_20_H_30_NO | 300.2324 | 0.5 |
| C5:1-AQ | [M+H]^+^ | C_14_H_16_NO | 214.1226 | -0.3 |
| C6:1-AQ | [M+H]^+^ | C_15_H_18_NO | 228.1382 | -0.3 |
| C7:1-AQ | [M+H]^+^ | C_16_H_20_NO | 242.1538 | -0.4 |
| C8:1-AQ | [M+H]^+^ | C_17_H_22_NO | 256.1690 | -2.4 |
| C9:1-AQ | [M+H]^+^ | C_18_H_24_NO | 270.1852 | -0.1 |
| C10:1-AQ | [M+H]^+^ | C_19_H_26_NO | 284.2004 | -1.6 |
| C11:1-AQ | [M+H]^+^ | C_20_H_28_NO | 298.2166 | 0.1 |
| C12:1-AQ | [M+H]^+^ | C_21_H_30_NO | 312.2310 | -3.9 |
| C13:1-AQ | [M+H]^+^ | C_22_H_32_NO | 326.2482 | 1.0 |
| C7-PQS/C7-NO (HQNO) | [M+H]^+^ | C_16_H_22_NO_2_ | 260.1645 | 0.1 |
| C8-PQS/C8-NO | [M+H]^+^ | C_17_H_24_NO_2_ | 274.1806 | 1.5 |
| C9-PQS/C9-NO (NQNO) | [M+H]^+^ | C_18_H_26_NO_2_ | 288.1958 | 0.0 |
| C11-PQS/C11-NO | [M+H]^+^ | C_20_H_30_NO_2_ | 316.2272 | 0.2 |
| C5-NO | [M+H]^+^ | C_14_H_18_NO_2_ | 232.1332 | 0.1 |
| C6-NO | [M+H]^+^ | C_15_H_20_NO_2_ | 246.1490 | 0.6 |
| C10-NO | [M+H]^+^ | C_19_H_28_NO_2_ | 302.2113 | -0.4 |
| C7:1-NO | [M+H]^+^ | C_16_H_20_NO_2_ | 258.1490 | 0.4 |
| C8:1-NO | [M+H]^+^ | C_17_H_22_NO_2_ | 272.1643 | -0.9 |
| C9:1-NO | [M+H]^+^ | C_18_H_24_NO_2_ | 286.1802 | 0.0 |
| C10:1-NO | [M+H]^+^ | C_19_H_26_NO_2_ | 300.1964 | 1.9 |
| C11:1-NO | [M+H]^+^ | C_20_H_28_NO_2_ | 314.2115 | 0.0 |
| C12:1-NO | [M+H]^+^ | C_21_H_30_NO_2_ | 328.2269 | -0.8 |
| **3-alkyl-2,3-dihydroxy-4-quinolones** | | | | |
| C7 | [M+H]^+^ | C_16_H_22_NO_3_ | 276.1594 | -0.1 |
| C9 | [M+H]^+^ | C_18_H_26_NO_3_ | 304.1908 | 0.2 |
| C7:1 | [M+H]^+^ | C_16_H_20_NO_3_ | 274.1437 | -0.2 |
| C9:1 | [M+H]^+^ | C_18_H_24_NO_3_ | 302.1751 | 0.3 |
| **Rhamnolipids** | | | | |
| Rha-C10 | [M+NH_4_]^+^ | C_16_H_34_NO_7_ | 352.2322 | -2.3 |
| Rha-C10-C10 | [M+NH_4_]^+^ | C_26_H_52_NO_9_ | 522.3639 | 0.5 |
| Rha-C12:1-C10 | [M+NH_4_]^+^ | C_28_H_54_NO_9_ | 548.3784 | -1.6 |
| Rha-C10-C12/C12-C10 | [M+NH_4_]^+^ | C_28_H_56_NO_9_ | 550.3950 | 0.1 |
| Rha-Rha-C10-C10 | [M+NH_4_]^+^ | C_32_H_62_NO_13_ | 668.4214 | -0.3 |
| Rha-Rha-C10-C12/C12-C10 | [M+NH_4_]^+^ | C_34_H_66_NO_13_ | 696.4530 | 0.1 |
| Rha-Rha-C10 | [M+NH_4_]^+^ | C_22_H_44_NO_11_ | 498.2910 | 0.3 |
| ***N*-acyl-homoserine lactone (AHLs)** | | | | |
| C4-AHL | [M+H]^+^ | C_8_H_14_NO_3_ | 172.0967 | -0.9 |
| C6-AHL | [M+H]^+^ | C_10_H_18_NO_3_ | 200.1281 | -0.3 |
| 3-Oxo-C4-AHL | [M+H]^+^ | C_8_H_12_NO_4_ | 186.0760 | -0.4 |
| 3-OH-C4-AHL | [M+H]^+^ | C_8_H_14_NO_4_ | 188.0916 | -0.5 |
| 3-Oxo-C6-AHL | [M+H]^+^ | C_10_H_16_NO_4_ | 214.1071 | -0.7 |
| 3-OH-C6-AHL | [M+H]^+^ | C_10_H_18_NO_4_ | 216.1230 | -0.3 |
| **Nucleobases** | | | | |
| thymine | [M+H]^+^ | C_5_H_7_N_2_O_2_ | 127.0502 | 0.0 |
| Uracil | [M+H]^+^ | C_4_H_5_N_2_O_2_ | 113.0345 | -0.1 |
| Cytosine | [M+H]^+^ | C_4_H_6_N_3_O | 112.0505 | 0.0 |
| Adenine | [M+H]^+^ | C_5_H_6_N_5_ | 136.0617 | -0.5 |
| Guanine | [M+H]^+^ | C_5_H_6_N_5_O | 152.0566 | -0.8 |
| **Amino Acids** | | | | |
| Arginine | [M+H]^+^ | C_6_H_15_N_4_O_2_ | 175.1189 | -0.6 |
| Valine | [M+H]^+^ | C_5_H_12_NO_2_ | 118.0862 | -0.4 |
| Glutamine | [M+H]^+^ | C_5_H_11_N_2_O_3_ | 147.0763 | -0.9 |
| Serine | [M+H]^+^ | C_3_H_8_NO_3_ | 106.0499 | -0.3 |
| Aspartic acid | [M+H]^+^ | C_4_H_8_NO_4_ | 134.0446 | -1.0 |
| L-Threonine/Homoserine | [M+H]^+^ | C_4_H_10_NO_3_ | 120.0654 | -0.5 |
| cysteine | [M+H]^+^ | C_3_H_8_NO_2_S | 122.0270 | -0.9 |
| homocysteine | [M+H]^+^ | C_4_H_10_NO_2_S | 136.0426 | -1.2 |
| methionine | [M+H]^+^ | C_5_H_12_NO_2_S | 150.0582 | -1.0 |
| Isoleucine | [M+H]^+^ | C_6_H_14_NO_2_ | 132.1017 | -0.9 |
| lysine | [M+H]^+^ | C_6_H_15_N_2_O_2_ | 147.1125 | -1.2 |
| Glutamic acid | [M+H]^+^ | C_5_H_10_NO_4_ | 148.0604 | -1.1 |
| ornithine | [M+H]^+^ | C_5_H_13_N_2_O_2_ | 133.0972 | -1.0 |
| N-α-Acetylcitrulline | [M+H]^+^ | C_8_H_16_N_3_O_4_ | 218.1131 | -0.4 |
| Proline | [M+H]^+^ | C_5_H_10_NO_2_ | 116.0706 | -0.8 |
| histidine | [M+H]^+^ | C_6_H_10_N_3_O_2_ | 156.0764 | -0.8 |
| L-tryptophan | [M+H]^+^ | C_11_H_13_N_2_O_2_ | 205.0971 | -0.5 |
| L-phenylalanine | [M+H]^+^ | C_9_H_12_NO_2_ | 166.0862 | -1.0 |
| tyrosine | [M+H]^+^ | C_9_H_12_NO_3_ | 182.0811 | -0.4 |
| Pretyrosine | [M+H]^+^ | C_10_H_14_NO_5_ | 228.0873 | -0.4 |
| **Lipids** | | | | |
| PE(15:0cyclo/19:0cycv8c) | [M+H]^+^ | C_39_H_75_NO_8_P | 716.5225 | 0.0 |
| PE(19:0cycv8c/15:0cyclo) |  |  |  |  |
| PE(16:1(9Z)/18:1(11Z)) |  |  |  |  |
| PE(18:1(11Z)/16:1(9Z)) |  |  |  |  |
| PE(17:0cycw7c/17:0cycw7c) |  |  |  |  |
| PE(16:1(9Z)/18:1(9Z)) |  |  |  |  |
| PE(18:1(9Z)/16:1(9Z)) |  |  |  |  |
| PE(18:1(11Z)/16:0) | [M+H]^+^ | C_39_H_77_NO_8_P | 718.5380 | -0.1 |
| PE(16:0/18:1(11Z)) | [M+K]^+^ | C_39_H_76_NO_8_PK | 756.4938 | -0.3 |
| PE(15:0cyclo/19:iso) | [M+Na]^+^ | C_39_H_76_NO_8_PNa | 740.5200 | -0.2 |
| PE(19:iso/15:0cyclo) |  |  |  |  |
| PE(16:0/18:1(9Z)) |  |  |  |  |
| PE(16:1(9Z)/18:0) |  |  |  |  |
| PE(18:0/16:1(9Z)) |  |  |  |  |
| PE(18:1(9Z)/16:0) |  |  |  |  |
| **Miscellaneous compounds** |  |  |  |  |
| dihydroxyquinolone (DHQ) | [M+H]^+^ | C_9_H_8_NO_2_ | 162.0548 | -1.1 |
| 2-Aminoacetophenone | [M+H]^+^ | C_8_H_10_NO | 136.0756 | -0.5 |
| Kynurenine | [M+H]^+^ | C_10_H_13_N_2_O_3_ | 209.0921 | 0.4 |
| Anthranilic acid | [M+H]^+^ | C_7_H_8_NO_2_ | 138.0548 | -0.8 |
| Palmitic Acid (C16:0) | [M+NH_4_]^+^ | C_16_H_36_NO_2_ | 274.2741 | 0.2 |
| Glutathione | [M+H]^+^ | C_10_H_18_N_3_O_6_S | 308.0911 | 0.0 |
| AMP/dGMP | [M+H]^+^ | C_10_H_13_N_5_O_7_P | 348.0705 | 0.3 |
| cis-2-decenoic acid (CDA) | [M+NH_4_]^+^ | C_10_H_22_NO_2_+ | 188.1642 | -1.4 |
| ectoine | [M+H]^+^ | C_6_H_11_N_2_O_2_+ | 143.0815 | -0.2 |
| 2-oxobutanoate | [M+H]^+^ | C_4_H_7_O_3_ | 103.0390 | -0.2 |
| 2-oxoisovalerate | [M+H]^+^ | C_5_H_9_O_3_ | 117.0546 | -0.4 |
| 2-Aminohexanedioic acid | [M+H]^+^ | C_6_H_12_NO_4_ | 162.0758 | -1.9 |
| phosphoenolpyruvate | [M+H]^+^ | C_3_H_6_O_6_P | 168.9896 | -0.6 |
| D-erythrose-4-phosphate | [M+H]^+^ | C_4_H_10_O_7_P | 201.0160 | 0.5 |
| 2-Oxoadipic acid | [M+H]^+^ | C_6_H_9_O_5_ | 161.0444 | -0.4 |

**Supplementary Table 2.** Secondary ion intensities of compounds from frozen-hydrated biofilm and freeze-dried biofilm in GCIB-Orbitrap spectra and the relative intensity ratio and XLogP where available. The value with * in the column of Peak Area 1 means the signal intensity is too low and this value is noise; The compound with ^≠^ in the column of Assignment means the adduct is [M+NH_4_]^+^ or [M+Na]^+^ or [M+K]^+^ instead of [M+H]^+^. XLogP data are from PubChem. The data with ^×^ in the column of XLogP come from PAMDB database. XLogP filled with N/A is either because the LogP of this compound cannot be found or the adduct of this compound is not [M+H]^+^ which were not considered in **Supplementary Figure 18.**

| *m/z* | Chemical Formula | Assignment | Peak Area 1 (freeze-dried samples) | Peak Area 2 (frozen-hydrated samples) | Ratio of peak area 2 to peak area 1 | XLogP |
| --- | --- | --- | --- | --- | --- | --- |
| **Amino acids** | | | | | | |
| 106.0499 | C_3_H_8_NO_3_ | Serine | *2973 | 82265887 | 27673 | -3.1 |
| 134.0446 | C_3_H_8_NO_3_ | Aspartic acid | *33972 | 44141541 | 1299 | -5.6 |
| 120.0654 | C_3_H_10_NO_3_ | L-threonine/  Homoserine | *2799 | 135024754 | 48246 | -2.9 |
| 122.0270 | C_3_H_8_NO_2_S | Cysteine | *2803 | 154214 | 55 | -2.5 |
| 136.0426 | C_3_H_10_NO_2_S | Homocysteine | *2864 | 144008 | 50 | -3.4 |
| 150.0582 | C_5_H_12_NO_2_S | Methionine | *3310 | 1033428 | 312 | -1.9 |
| 132.1017 | C_6_H_13_NO_2_ | Isoleucine | *3514 | 71421242 | 20322 | -1.7 |
| 147.1125 | C_6_H_15_N_2_O_2_ | lysine | *2029 | 193532534 | 95392 | -3.0 |
| 148.0604 | C_5_H_10_NO_3_ | Glutamic acid | *3450 | 253199567 | 73402 | -3.7 |
| 133.0972 | C_5_H_13_N_2_O_2_ | Ornithine | *2587 | 137516558 | 53149 | -4.4 |
| 218.1131 | C_8_H_16_N_3_O_3_ | N- α -Acetylcitrulline | *3656 | 11989788 | 3280 | -2.3 |
| 116.0706 | C_5_H_10_NO_2_ | Proline | 13328 | 157826170 | 11842 | -2.5 |
| 156.0764 | C_6_H_10_N_3_O_2_ | Histidine | 21009 | 38991110 | 1856 | -3.2 |
| 175.1189 | C_6_H_15_N_3_O_2_ | Arginine | *7752 | 31513155 | 4065 | -4.2 |
| 205.0971 | C_11_H_13_N_2_O_2_ | L-tryptophan | 11753 | 2717816 | 231 | -1.1 |
| 166.0862 | C_9_H_12_NO_2_ | L-phenylalanine | 25876 | 13655804 | 528 | -1.5 |
| 182.0811 | C_9_H_12_NO_3_ | Tyrosine | *6732 | 20691398 | 3074 | -2.3 |
| 228.0873 | C_10_H_13_NO_5_ | Pretyrosine | *3683 | 286064 | 78 | -3.1 |
| 118.0862 | C_5_H_12_NO_2_ | Valine | 63436 | 117977101 | 1860 | -2.3 |
| 147.0761 | C_5_H_11_N_2_O_3_ | Glutamine | *3455 | 93543636 | 27076 | -3.1 |
|  |  |  |  | Average ratio | 18690 |  |
| **Nucleobases** | | |  |  |  |  |
| 127.0502 | C_5_H_7_N_2_O_2_ | Thymine | 120617 | 70229225 | 582 | -0.6 |
| 113.0345 | C_4_H_5_N_2_O_2_ | Uracil | 134784 | 58826006 | 436 | -1.1 |
| 112.0505 | C_4_H_6_N_3_O | cytosine | 25645145 | 1236343235 | 48 | -1.7 |
| 136.0618 | C_5_H_6_N_5_ | Adenine | 61033669 | 977882528 | 16 | -0.1 |
| 152.0566 | C_5_H_6_N_5_O | Guanine | 21692565 | 926216200 | 43 | -1.1 |
|  |  |  |  | Average ratio | 225 |  |
| **Alkyl quinolones** | | | | | | |
| 216.1383 | C_14_H_18_NO | C5-AQ | 18198 | 335683 | 18 | 3.3 |
| 230.1539 | C_15_H_20_NO | C6-AQ | 11702 | 103578 | 9 | ^×^4.6 |
| 244.1696 | C_16_H_22_NO | C7-AQ (HHQ) | 2006280 | 19242473 | 10 | 4.9 |
| 258.1852 | C_17_H_24_NO | C8-AQ | 27407 | 175216 | 6 | ^×^5.4 |
| 272.2008 | C_18_H_26_NO | C9-AQ (NHQ) | 12961080 | 155214959 | 12 | 6.0 |
| 286.2165 | C_19_H_28_NO | C10-AQ | 83359.63 | 115171 | 1 | ^×^6.3 |
| 300.2322 | C_20_H_30_NO | C11-AQ | 320420.96 | 5689358 | 18 | 6.5 |
| 214.1226 | C_14_H_16_NO | C5:1-AQ | 58613.21 | 943016 | 16 | N/A |
| 228.1383 | C_15_H_18_NO | C6:1-AQ | 21833.12 | 464892 | 21 | N/A |
| 242.1539 | C_16_H_20_NO | C7:1-AQ | 55862.49 | 2251481 | 40 | N/A |
| 256.1695 | C_17_H_22_NO | C8:1-AQ | 13132.01 | 133024 | 10 | N/A |
| 270.1852 | C_18_H_24_NO | C9:1-AQ | 3464370.47 | 103766170 | 30 | N/A |
| 284.2009 | C_19_H_26_NO | C10:1-AQ | 27591.56 | 269971 | 10 | N/A |
| 298.2166 | C_20_H_28_NO | C11:1-AQ | 595039 | 30772443 | 52 | N/A |
| 312.2321 | C_21_H_30_NO | C12:1-AQ | 14184 | 82672 | 6 | N/A |
| 326.2479 | C_22_H_32_NO | C13:1-AQ | 19005 | 537583 | 28 | N/A |
| 260.1645 | C_16_H_22_NO_2_ | C7-PQS/C7-NO (HQNO) | 1359102 | 81805809 | 60 | 4.5 |
| 274.1800 | C_17_H_24_NO_2_ | C8-PQS/C8-NO | *5383 | 319812 | 59 | ^×^4.1 |
| 288.1958 | C_18_H_26_NO_2_ | C9-PQS/C9-NO (NQNO) | 1867845 | 62479058 | 33 | 4.6 |
| 316.2271 | C_20_H_30_NO_2_ | C11-PQS/C11-NO | 8479 | 622327 | 73 | ^×^5.5 |
| 232.1331 | C_14_H_18_NO_2_ | C5-NO | *4496 | 370778 | 82 | ^×^2.7 |
| 246.1489 | C_15_H_20_NO_2_ | C6-NO | *3897 | 157108 | 40 | ^×^3.2 |
| 302.2111 | C_19_H_28_NO_2_ | C10-NO | *4266 | 124468 | 29 | ^×^5.0 |
| 258.1489 | C_16_H_20_NO_2_ | C7:1-NO | 9637 | 5941676 | 617 | ^×^3.2 |
| 272.1644 | C_17_H_22_NO_2_ | C8:1-NO | *2741 | 132773 | 48 | N/A |
| 286.1802 | C_18_H_24_NO_2_ | C9:1-NO | 664974 | 44125738 | 66 | ^×^4.1 |
| 300.1955 | C_19_H_26_NO_2_ | C10:1-NO | *5686 | 208303 | 37 | N/A |
| 314.2114 | C_20_H_28_NO_2_ | C11:1-NO | 191149 | 11326083 | 59 | ^×^5.0 |
| 328.2271 | C_21_H_30_NO_2_ | C12:1-NO | *3818 | 38351 | 10 | N/A |
| 276.1593 | C_16_H_22_NO_3_ | C7 | *4888 | 1771830 | 362 | ^×^2.6 |
| 304.1902 | C_18_H_26_NO_3_ | C9 | *2369 | 717805 | 303 | ^×^3.4 |
| 274.1439 | C_16_H_20_NO_3_ | C7:1 | *3582 | 123447 | 34 | ^×^2.4 |
| 302.1752 | C_18_H_24_NO_3_ | C9:1 | *2744 | 371070 | 135 | ^×^3.3 |
|  |  |  |  | Average ratio | 71 |  |
| ***N*-acyl homoserine lactones** |  |  |  |  |  |  |
| 172.0967 | C_8_H_14_NO_3_ | C4-AHL | *3040 | 14704066 | 4837 | 0.5 |
| 200.1277 | C_10_H_18_NO_3_ | C6-AHL | *1952 | 2102223 | 1077 | 1.5 |
| 186.0768 | C_8_H_12_NO_4_ | 3-OxO-C4-AHL | *13021 | 17552822 | 1348 |  |
| 214.1071 | C_10_H_16_NO_4_ | 3-Oxo-C6-AHL | *2709 | 1072811 | 396 | 0.9 |
| 188.0916 | C_8_H_14_NO_4_ | 3-OH-C4-AHL | *3333 | 56656298 | 16999 | -0.1 |
| 216.1236 | C_10_H_18_NO_4_ | 3-OH-C6-AHL | *2538 | 797007 | 314 | N/A |
|  |  |  |  | Average ratio | 4162 |  |
| **Rhamnolipids** | | | | | | |
| 352.2331 | C_16_H_34_NO_7_ | ^≠^Rha-C10 | *5059 | 247838 | 49 | N/A |
| 522.3639 | C_26_H_52_NO_9_ | ^≠^Rha-C10-C10 | *2896 | 1764452 | 609 | N/A |
| 548.3794 | C_28_H_54_NO_9_ | ^≠^Rha-C12:1-C10 | *2919 | 124049 | 42 | N/A |
| 550.3947 | C_28_H_56_NO_9_ | ^≠^Rha-C10-C12/C12-C10 | *4095 | 189577 | 46 | N/A |
| 668.4216 | C_32_H_62_NO_13_ | ^≠^Rha-Rha-C10-C10 | *2810 | 611414 | 218 | N/A |
| 696.4527 | C_34_H_66_NO_13_ | ^≠^Rha-Rha-C10-C12/C12-C10 | *3213 | 4499917 | 1401 | N/A |
| 498.2918 | C_22_H_44_NO_11_ | ^≠^Rha-Rha-C10 | *3456 | 758975 | 220 | N/A |
|  |  |  |  | Average ratio | 369 |  |
| **Lipids** |  |  |  |  |  |  |
| 740.5200 | C_39_H_76_NO_8_PNa | ^≠^PE lipids | 21002 | 13478674 | 642 | N/A |
| 756.4939 | C_39_H_76_NO_8_PK | ^≠^PE lipids | 60737 | 2797571 | 46 | N/A |
| 716.5222 | C_39_H_75_NO_8_P | PE lipids | *3617 | 21167995 | 5853 | 7-12 |
| 718.5380 | C_39_H_77_NO_8_P | PE lipids | *5353 | 56546324 | 10564 | 7-12 |
| 142.0262 | C_2_H_9_NO_4_P | PE lipids head | *3441 | 317245000 | 92200 | N/A |
|  |  |  |  | Average ratio | 21861 |  |
| **Miscellaneous compounds** | |  |  |  |  |  |
| 162.0549 | C_9_H_8_NO_2_ | DHQ | 2137113 | 5928465 | 3 | 1.6 |
| 136.0757 | C_8_H_10_NO | 2-Aminoacetophenone | 20121683 | 243866840 | 12 | 0.8 |
| 209.0921 | C_10_H_13_N_2_O_3_ | Kynurenine | 8586 | 3122311 | 364 | -2.2 |
| 138.0549 | C_7_H_8_NO_2_ | Anthranilic acid | 1734388 | 36389056 | 21 | 1.2 |
| 274.2740 | C_16_H_36_NO_2_ | ^≠^Palmitic Acid (C16:0) | *1708 | 1098297 | 643 | N/A |
| 188.1642 | C_10_H_22_NO_2_ | ^≠^cis-2-decenoic acid (CDA) | *4520 | 3524071 | 780 | N/A |
| 181.0703 | C_6_H_13_O_6_ | galactose/mannose | *2203 | 3311541 | 1503 | -2.6 |
| 348.0688 | C_10_H_15_N_5_O_7_P | AMP or GMP | 17490 | 20370256 | 1165 | -3.5 |
| 143.0815 | C_6_H_11_N_2_O_2_ | ectoine | 40832 | 118663212 | 2906 | -1 |
| 89.0236 | C_3_H_5_O_3_ | Pyruvate | *2746 | 25196550 | 9177 | -0.6 |
| 103.0390 | C_4_H_7_O_3_ | 2-oxobutanoate | *2929 | 35245213 | 12032 | 0.1 |
| 117.0546 | C_5_H_9_O_3_ | 2-oxoisovalerate | *4573 | 24673743 | 5395 | 0.7 |
| 162.0758 | C_6_H_12_NO_4_ | 2-Aminohexanedioic acid | *3178 | 75426386 | 23734 | -3.1 |
| 168.9896 | C_3_H_6_O_6_P | phosphoenolpyruvate | *2482 | 361955 | 146 | -1.1 |
| 201.0160 | C_4_H_10_O_7_P | D-erythrose-4-phosphate | *3827 | 22002865 | 5749 | -3.3 |
| 308.0910 | C_10_H_18_N_3_O_6_S | Glutathione | *2973 | 66279324 | 22296 | -4.5 |
| 161.0444 | C_6_H_9_O_5_ | 2-Oxoadipic acid | *3149 | 14195306 | 4509 | -0.5 |
|  |  |  |  | Average ratio | 5320 |  |

**Supplementary Video 1.** z-Axis cropping of 3D rendered ToF MS images of the frozen hydrated biofilm with adenine/2-Aminoacetophenone (*m/z* 136) on the substrate (Al, *m/z* 27)

**Supplementary Video 2.** z-Axis cropping of 3D rendered ToF MS images of the frozen hydrated biofilm with K^+^ (*m/z* 39) on the substrate (Al, m*/z* 27, shown in grey)
